# A cyclin-polarity feedback network ensures healthy cell proliferation

**DOI:** 10.1101/2025.08.18.670832

**Authors:** Landry Peyran, Charles Lefranc, Steven P. Gygi, Anne Royou, Derek McCusker

**Affiliations:** University of Bordeaux, CNRS, UMR 5095; Bordeaux, Nouvelle Aquitaine, France; Department of Cell Biology, Harvard Medical School; Boston, MA, USA

## Abstract

Healthy proliferation requires the coordination of cell cycle progression with cell polarity. In budding yeast, polarity is established when G1-cyclin-Cdc28^Cdk1^ triggers Cdc42 activation to generate a cell pole that is used as an axis for growth and division. While polarity defects delay the cell cycle temporally, permitting error correction, it is unknown if Cdc28^Cdk1^ directly rectifies errant polarity. Here, we identify an adaptive response where G1-cyclin-Cdc28^Cdk1^ participates in error correction via the augmentation of its kinase activity towards substrates that activate Cdc42. The response involves temporal and spatial cell cycle reconfiguration via extended G1 cyclin expression, nucleocytoplasmic rerouting and signaling. However, this strategy has a cost: if the defect is irreparable, high G1-cyclin levels enforce inexorable cell cycle commitment in the absence of a daughter cell, generating multinucleate cells. G1-cyclins therefore not only trigger G1 events, but also monitor their execution, employing feedback to coordinate polarity with cell cycle progression.

---

The decision to commit to cell division involves the integration of growth signals within the cell cycle control system, leading to growth-dependent G1 cyclin synthesis. G1 events are perhaps best, though by no means exhaustively understood in the budding yeast *Saccharomyces cerevisiae*. Here, G1 cyclin-associated Cdc28^Cdk1^ triggers decisive cell cycle commitment at the G1/S transition ^1^. The irreversibility of cell cycle commitment is thought to be mediated by a positive feedback loop where the G1-cyclin Cln3-Cdc28^Cdk1^ phosphorylates and partially inhibits Whi5, a transcriptional repressor of the G1 cyclins *CLN1* and *CLN2* ^2–4^. When relieved of their repression, expression of Cln1 and Cln2 fully phosphorylate and inhibit Whi5, thereby stimulating more Cln1 and Cln2 expression and a burst of associated Cdc28^Cdk1^ activity ^5,6^. In addition to driving an extensive G1 transcriptional program, or regulon, G1 cyclin Cdc28^Cdk1^ kinase also feeds into cell polarity, which at this point in the cell cycle is in a low activity state as reflected by a depolarized organization of actin filaments. G1 cyclin-Cdc28^Cdk1^ ensures the activation of the Rho GTPase Cdc42 to a level compatible with rapid, stable polarity axis establishment and the polarization of the actin cytoskeleton, which supports the growth of a daughter cell called the bud ^7^. This involves the nuclear export of the guanine exchange factor (GEF) for Cdc42, called Cdc24, into the cytoplasm ^8,9^. Concomitantly, phosphorylation and inactivation of Cdc42 GTPase Activating Proteins (GAPs) including Rga2 occurs, together with phosphorylation of the GEF Cdc24 and Cdc42 scaffold-associated proteins, triggering polarity axis establishment ^10–13^.

Importantly, G1 cyclin-Cdc28^Cdk1^ activity displays more than 10-fold substrate specificity towards these proteins compared to M-phase cyclin-Cdc28^Cdk1^ *in vitro* ^11^. This substrate specificity, which is likely mediated via G1 cyclin docking motifs in substrates, could explain why budding in *S. cerevisiae* only occurs at G1/S phase in the cell cycle, and why it is not reinitiated subsequently as Cdc28^Cdk1^ activity rises during S-phase and mitosis ^14,15^. The specific targeting of G1 cyclin activity towards the activation of the Cdc42 GTPase module is also borne out *in vivo*. *S. cerevisiae* strains in which all S- and M-phase cyclins are deleted can proliferate if a single engineered M-phase cyclin is provided, indicating that M-phase cyclins can drive S-phase events. However, the engineered M-phase cyclin cannot compensate for the loss of G1 cyclin activity, as it fails to drive polarity axis establishment and budding, indicative of G1 cyclin specificity towards polarity proteins *in vivo* ^16^.

Early in G1, Cdc42 is localized on the plasma membrane due to prenylation and polybasic residues at its C-terminus ^17,18^. These sequences anchor the protein to the plasma membrane, while the activation of the GTPase by its GEF Cdc24, and subsequent stabilization by the scaffold Bem1 drive its anisotropic activation during polarity axis establishment when G1 cyclin-Cdc28^Cdk1^ levels rise ^19,20^. Cdc42 is not distributed homogeneously in the plane of the plasma membrane, but is rather clustered in diffraction-limited ensembles termed nanoclusters. Nanoclustering is a conserved feature of Rho GTPase organization in yeast, plants and mammals ^21–24^. Moreover, it extends to the wider Ras-superfamily, where models propose that nanoclustering may digitize GTPase signaling by providing a high local concentration of GTPase module components ^25^. During polarity establishment in budding yeast, anionic lipids including phosphatidylserine and phosphoinositols such as PI4P and PI(4,5)P2 are enriched in the interior leaflet of the plasma membrane where they become polarized ^26–28^. These lipid species have also been demonstrated to play an important role in the nanoclustering of Ras and Rho-GTPases ^22–24,29^. Multivalent electrostatic interactions between anionic lipids and basic residues in Bem1’s N-terminus and PX domain, together with the PH domain of Cdc24 are required for the recruitment of the proteins to the pole ^30^. Here, Bem1 stimulates Cdc24 GEF activity and the generation of Cdc42-GTP ^31,32^, which may trigger a positive feedback loop that recruits more Bem1 and Cdc24 ^19,33^. The outcome of this signaling is a sharp rise in the polarized activation of Cdc42, emanating from plasma membrane nanoclusters. Mutation of the basic residues in Bem1 and Cdc24 that mediate their interaction with anionic lipids results in a severe loss of the anisotropic organization of the GEF and scaffold from the plasma membrane at steady-state, together with defects in the size of Cdc42 nanoclusters and the ablation of PAK signaling ^30^. However, the contribution of these multivalent lipid interactions to the dynamics of cell polarity and to ensuing cell cycle control has not been studied, and is the focus of this work.

Aberrant cell polarity and polarized growth are monitored by a cell cycle checkpoint in *S. cerevisiae* ^34,35^. Defects in Cdc42 signaling or in actin organization early in the cell cycle result in Swe1^Wee1^ kinase activation, which phosphorylates and inhibits M-phase Cdc28^Cdk1 36^, thus delaying nuclear division ^37,38^. Swe1^Wee1^ activity interferes with both entry into mitosis and anaphase onset ^39–43^. While Swe1 activation may provide a temporal window for polarity correction, it is unknown if Cdc28^Cdk1^ activity participates in the correction of polarity defects. The requirement for G1 cyclin-Cdc28^Cdk1^ in Cdc42 activation during polarity establishment in wild type cells raised the possibility of its involvement in correcting polarity problems stemming from sub-critical Cdc42 activity.

In this work, the *bem1 cdc24* anionic lipid binding mutant was used to investigate how interfering with Cdc42 nanoclustering affects the dynamics of cell polarity, enabling the identification of an unanticipated feedback relationship between GTPase signaling and G1 cyclins that is required for healthy cell proliferation.

## Results

### Polarity axis maintenance requires the lipid tethering of a Cdc42 GEF & scaffold to the plasma membrane

To assess how anionic lipid tethering of the Cdc42 activators Cdc24 and Bem1 affect the dynamics of cell polarity, the exocyst subunit Sec3-GFP, whose localization depends on Cdc42 activity ^44^, was monitored by time-lapse spinning disk microscopy in wild type and in *bem1 cdc24^lip^* mutant cells. In this mutant, multivalent anionic lipid binding sites were mutated in the Cdc24 PH domain, in Bem1’s PX domain and in N-terminal basic clusters in Bem1 that were previously identified (Supplementary Fig. 1a) ^30,45^. Cells were imaged from cell separation of one cell cycle (denoted time 0) until they formed medium sized buds in the next cell cycle, which includes the period at the G1/S transition in which budding yeast establish the polarity axis used for growth and division (green shading in Fig. 1a). In wild type control cells, Sec3-GFP became polarized, delineating a single, stable polarity axis (Fig. 1a, upper panel and Supplementary Movie 1). In contrast, the time from cell separation to Sec3-GFP polarization was delayed in the *bem1 cdc24^lip^* mutant, indicative of a delay in polarity establishment (Fig. 1a, lower panel and Fig. 1b). In addition, the mutant exhibited a severe polarity maintenance defect that manifested itself as an unstable pole (38% of cells). In the example shown, the cell made two attempts at Sec3-GFP polarization before successfully budding. As a result, the time from cell separation to bud emergence, and from Sec3-GFP polarization to bud emergence was delayed in the mutant compared to wild type cells (Fig. 1c and Supplementary Fig. 1b, 1c and Supplementary Movie 2). Polarity establishment is a cytoplasmic event controlled by G1-cyclin Cdc28^Cdk1^ activity ^11,46^. In order to determine if nuclear events driven by G1-cyclin Cdc28^Cdk1^ activity were also delayed in the mutant, we imaged Whi5-sfGFP, a transcriptional repressor acting at the G1/S transition whose nuclear export requires G1-cyclin Cdc28^Cdk1^ activity in a manner analogous to retinoblastoma (Rb) in metazoans ^3^. Strikingly, Whi5-sfGFP nuclear export occurred more rapidly in a synchronized population of *bem1 cdc24^lip^* mutant cells compared to wild type cells, suggesting a loss of coordination between nuclear and cytoplasmic G1-cyclin Cdc28^Cdk1^–driven events in the polarity mutant (Fig. 1d, e).

**Fig. 1.**
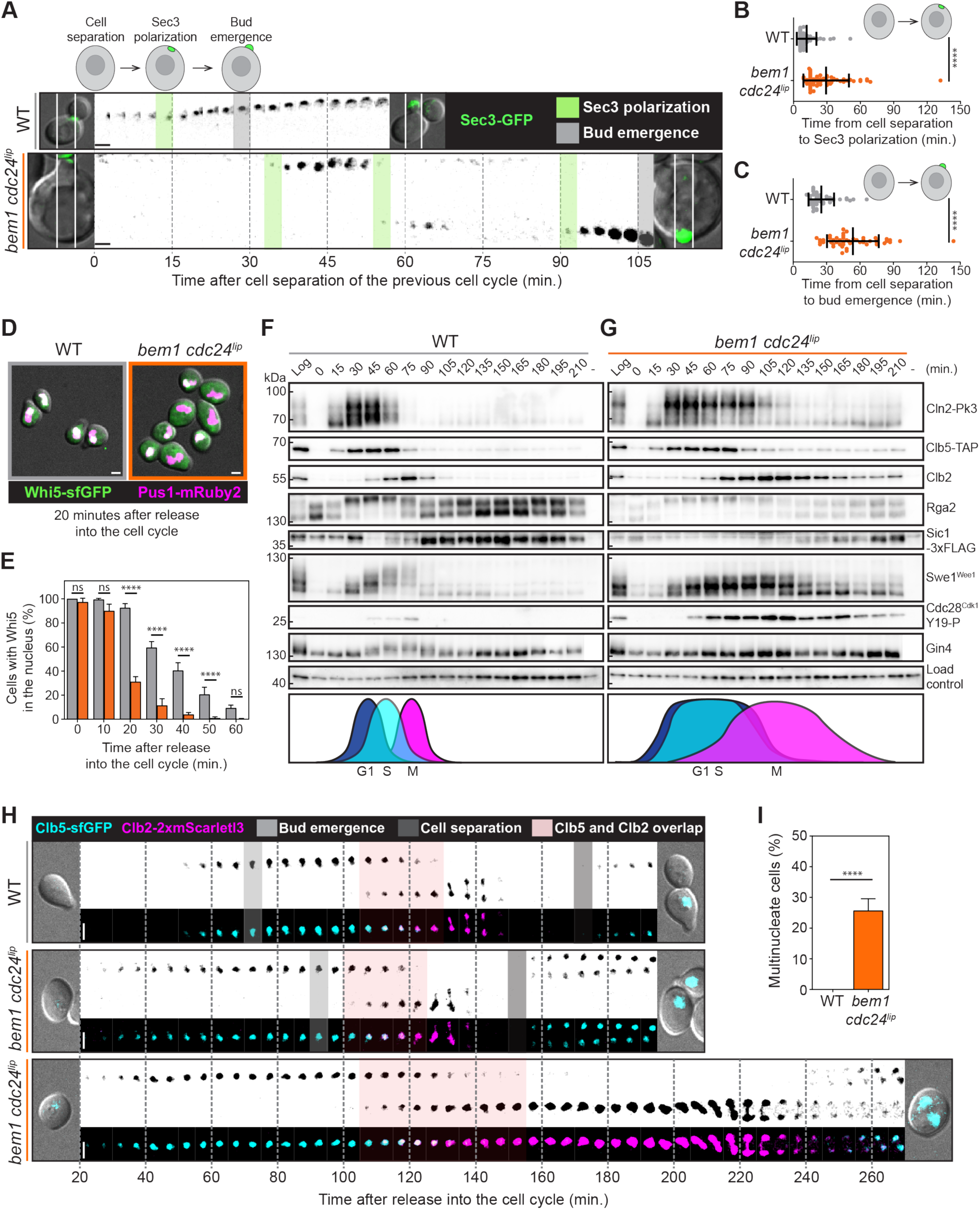
Lipid tethering defects in the *bem1 cdc24^lip^* mutant impair polarity maintenance and the coordination of nuclear and cytoplasmic events. **(a)** Exocyst (Sec3-GFP) dynamics in wild type and *bem1 cdc24^lip^* cells. Images are maximum-intensity projected z-stacks acquired every 3-minutes. Time 0 is cell separation in the previous cell cycle. **(b)** Time between cell separation and Sec3-GFP polarization in wild type (*n* = 53 cells) and *bem1 cdc24^lip^* (*n* = 61 cells). **(c)** Time between cell separation and bud emergence in wild type (*n* = 52 cells) and *bem1 cdc24^lip^* (*n* = 49 cells). **(d)** Whi5-sfGFP and Pus1-mRuby2 (nucleoplasm) 20 minutes after cell cycle entry. **(e)** Quantification of Whi5-sfGFP nuclear export. (*n* > 150 per time point). **(f)** Cell cycle progression after G1 synchronization in wild type cells. **(g)** Cell cycle in *bem1 cdc24^lip^*cells. **(h)** Dynamics of the S-phase cyclin Clb5-sfGFP (top, cyan), M-phase cyclin Clb2-2xmScarletl3 (middle, magenta) and both (bottom). An example of a cell that buds (middle) and that does not bud (bottom) is shown for the *bem1 cdc24^lip^* mutant. **(i)** Frequency of multinucleate cells in wild type (*n* = 684 cells) and the *bem1 cdc24^lip^* mutant (*n* = 945 cells). Values display mean +/-SD. Unpaired *t-*tests with Welch’s correction were performed in B, C and I (\*\*\*\**P* < 0.0001). A two-way ANOVA was performed in E (\*\*\*\**P* < 0.0001). Scale bars = 2 µm.

In order to understand how nuclear and cytoplasmic events become uncoordinated in the *bem1 cdc24^lip^* mutant, progression through a single cell cycle was monitored in cells synchronized in G1 with alpha-factor (Supplementary Fig. 1d). The cell cycle in control cells was characterized by successive temporal waves of cyclin-Cdc28^Cdk1^ activity generated by rapid increases and subsequent decline in the cyclins Cln2 (G1), Clb5 (S-phase) and Clb2 (M-phase) (Fig. 1f). Abrupt Cln2 cyclin synthesis drives decisive polarity establishment and bud emergence (Supplementary Fig. 1e), via phosphorylation of G1 cyclin substrates including the Cdc42 GAP Rga2 (Fig. 1f). The characteristic switch-like properties of the cell cycle are reinforced by the rapid inhibition of Cdc28^Cdk1^ inhibitors Sic1, whose degradation is initiated at the G1/S transition in response to G1-cyclin Cdc28^Cdk1^ activity ^47,48^, and Swe1, which undergoes multisite phosphorylation and inhibition at M-phase entry ^49^. These mechanisms form part of a wider control network ensuring that cells enter mitosis decisively with low levels of inhibitory Cdc28^Cdk1^ Y19 phosphorylation and mitosis-specific phospho-regulation of proteins such as Gin4 ^50^.

In contrast, bud emergence was less switch-like in *bem1 cdc24^lip^* cells (Supplementary Fig. 1e), where live-imaging had shown the cells undergoing repeated attempts at polarity establishment. The G1 cyclin Cln2 exhibited an extended temporal window of expression (Cln2 started to decline after 45 minutes in control cells and after 90 minutes in *bem1 cdc24^lip^* cells, Fig. 1g) and exhibited rapid and quantitative hyperphosphorylation. Consistent with the extended window of Cln2 expression, a longer period of Rga2 phosphorylation and low Sic1 expression was observed. Both events require Cln-Cdc28^Cdk1^ activity, indicating that phosphorylated G1 cyclins observed in the mutant are active. All of the cyclins examined, Cln2, Clb5 and Clb2, displayed sluggish dynamics in the mutant, suggesting that the cell cycle’s switch-like properties were compromised. Moreover, three lines of evidence were indicative of Swe1^Wee1^-kinase activation in the mutant: (1) high levels of inhibitory Cdc28^Cdk1^ Y19 phosphorylation; (2) an intermediate phosphorylated species of Swe1^Wee1^ that has previously been identified as the active form of the kinase ^49^, (3) the absence of mitosis-specific regulation of Gin4 ^51^.

To determine the consequences of Swe1^Wee1^ kinase activation on subsequent cell cycle progression, we monitored Clb5-sfGFP (S-phase) and Clb2-2xmScarlet-l3 (M-phase) cyclin dynamics during a cell cycle in wild type and *bem1 cdc24^lip^* mutant cells by dual-colour live-cell imaging (Fig. 1h). The single-cell data supported the conclusion from western blots that the expression of S- and M-phase cyclins were confined to a shorter, more discrete temporal window in wild type cells (Supplementary Movie 3), in contrast to the *bem1 cdc24^lip^* mutant, where loss of the cell cycle’s switch-like properties resulted in longer periods of cyclin expression (Fig. 1h and Supplementary Fig. 1f and g), and more extensive overlap of S- and M-phase cyclins (Fig. 1h and Supplementary Fig. 1h). Unbudded *bem1 cdc24^lip^* mutant cells displayed prolonged Clb2-2xmScarletl3 expression during an extended mitotic delay preceding anaphase (Fig. 1h, Supplementary Movies 4 and 5), consistent with elevated Swe1^Wee1^ kinase activity and high inhibitory Cdc28^Cdk1^ Y19 phosphorylation that was observed by western blotting. While Swe1^Wee1^-dependent Cdc28^Cdk1^ inhibition provided a temporal window for polarity correction, the transient nature of the delay meant that cells eventually underwent anaphase even if a daughter cell had not been generated. As a result, approximately 25% *bem1 cdc24^lip^* mutant cells were multinucleate (Fig. 1i and Supplementary Movie 6).

In summary, the firm tethering of the Cdc42 regulators Bem1 and Cdc24 to anionic lipids resolves Cdc42 activation into a stable pole that is required for timely polarity establishment and maintenance. Compromising lipid binding results in an increased window of active G1 cyclin expression during the cell cycle and an accompanying loss of coordination between nuclear and cytoplasmic cell cycle events driven by G1 cyclin Cdc28^Cdk1^ activity. When all recently identified anionic lipid binding sites in Bem1 and Cdc24 were mutated, the resulting polarity defects also triggered a Swe1^Wee1^-dependent mitotic delay via inhibitory phosphorylation of Cdc28^Cdk1^ on Y19 (Supplementary Fig. 1i). These results underscore the importance of multivalent plasma membrane tethering of Cdc42 activators for normal cell cycle progression.

### The severity of the polarity defect governs the extent of the cell cycle delay

The *bem1 cdc24^lip^* mutant had undergone generations of cell cycle progression during which the Swe1^Wee1^ checkpoint was active, raising the question of whether the observed phenotypes resulted from lipid binding defects, cell cycle checkpoint activation, or adaptation to sustained checkpoint activity (Supplementary Fig. 2a). A conditional system was therefore implemented to distinguish these possibilities. Bem1 was depleted using an auxin-inducible degron (AID) in *cdc24^lip^* cells co-transformed with integrated versions of WT *BEM1* or *bem1^lip^* (Fig. 2a)^52,53^. Having verified that the conditional system provided temporal control of bem1-AID depletion (Supplementary Fig. 2b-e), the AID system was used to study the immediate consequences of loss of Bem1 and Cdc24 lipid binding.

**Fig. 2.**
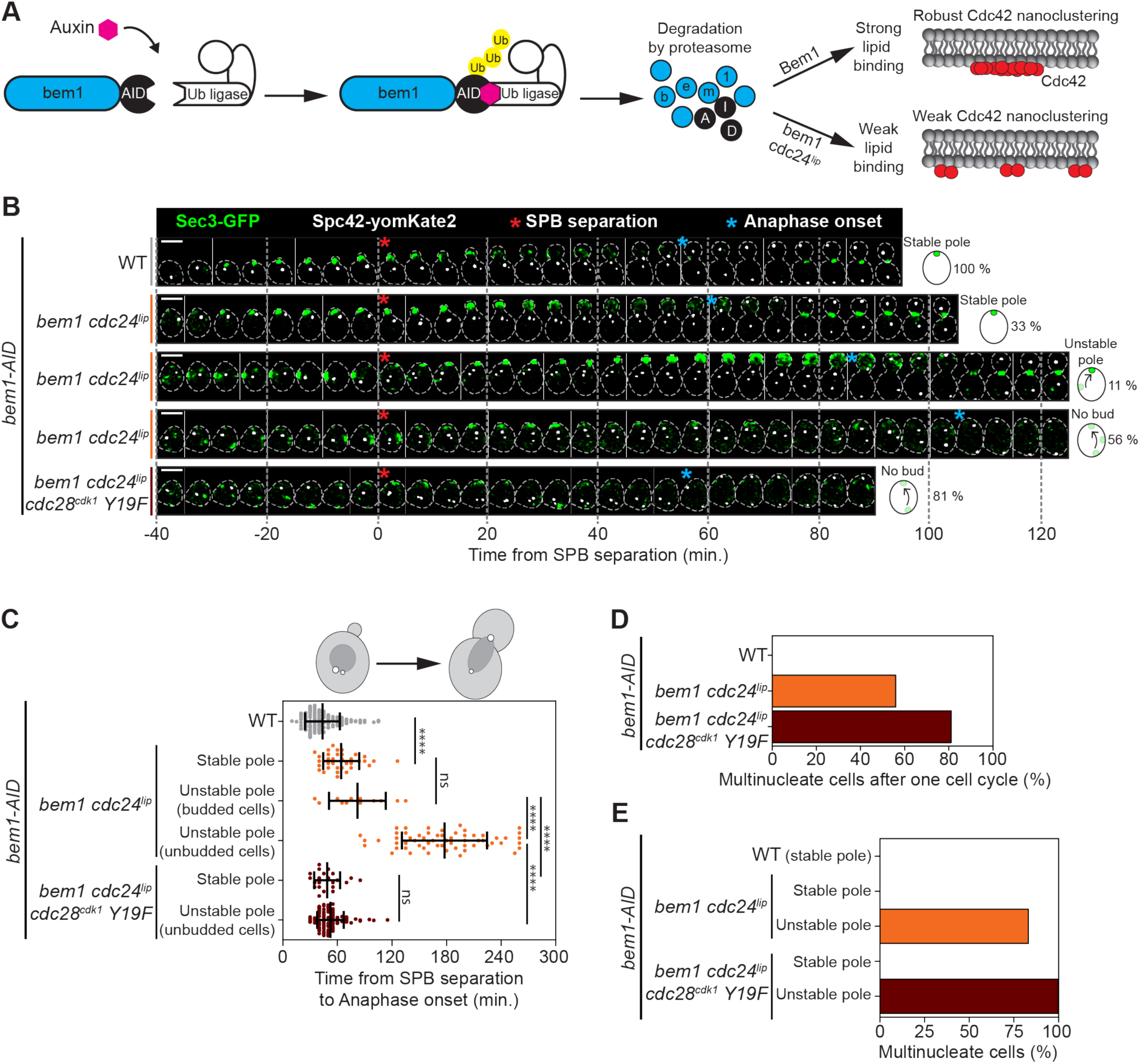
A conditional system indicates that the strength of the polarity defect in the *bem1 cdc24^lip^* mutant governs the extent of the Swe1^Wee1^ cell cycle delay. **(a)** Schematic of the AID system. **(b)** Exocyst (Sec3-GFP, green) and SPB (Spc42-yomKate2, white) dynamics after G1-arrest where auxin was added for the last 30-minutes of arrest. Time 0 indicates SPB separation. Images are maximum-intensity projected z-stacks displaying SPB separation (red asterisk) and anaphase (blue asterisk). Scale bars = 5 µm. **(c)** Time between SPB separation and anaphase for WT (*n* = 179), *bem1 cdc24^lip^* cells with stable Sec3 polarization (*n* = 35), unstable polarization that bud (*n* = 12), or unbudded (*n* = 60). Timing is also displayed for *bem1 cdc24^lip^ cdc28^cdk^*^1^ *Y19F* cells with stable (*n* = 22 cells) and unstable polarization (*n* = 95 cells). Values display mean +/- SD. Unpaired *t-*tests with Welch’s correction (\*\*\*\**P* < 0.0001). **(d)** Frequency of multinucleate cells in WT (*n* = 179), *bem1 cdc24^lip^* (*n* = 107) and *bem1 cdc24^lip^ cdc28^cdk^*^1^ Y19F cells (*n* = 117). **(e)** Frequency of WT cells (stable pole), *bem1 cdc24^lip^* cells exhibiting a stable (*n* = 35), or unstable pole (*n* = 72) that become multinucleate. The frequency of *bem1 cdc24^lip^ cdc28^cdk^*^1^ *Y19F* cells exhibiting stable (*n* = 22 cells) or unstable Sec3 polarization (*n* = 95 cells) that become multinucleate are indicated.

The dynamics of cell polarity and cell cycle progression were monitored simultaneously after bem1-AID depletion in *BEM1* cells or the *bem1 cdc24^lip^* mutant. The exocyst subunit Sec3-GFP was used as a marker of cell polarity, while Spc42-yomKate2, a spindle pole body component, was used as an indicator of cell cycle progression. Cells were synchronized in G1 and 1-NAA was added to degrade bem1-AID-6xFLAG 30-minutes prior to release into the cell cycle. Control cells entered the cell cycle with non-polarized Sec3-GFP that clustered as cells established a polarity axis. As cells budded, the Sec3-GFP signal was maintained in the growing bud (Fig. 2b). While 33**%** *bem1 cdc24^lip^*mutant cells established a stable pole of Sec3-GFP similarly to control cells, albeit at a slower rate (Fig. 2b and Supplementary Fig. 3a, stable pole), the remaining cells either formed an unstable pole that was then corrected, enabling cells to bud (unstable pole), or formed an unstable pole that was not corrected, preventing budding (no bud). Thus, the mutant displayed either a weak, moderate or strong polarity defect.

The panel of polarity phenotypes provided an opportunity to relate the severity of the polarity defects to the temporal cell cycle delay, which could not be addressed previously with actin polymerization toxins in which F-actin organization is ablated ^37,54^. The time from Sec3-GFP polarization to SPB separation was similar in the *bem1 cdc24^lip^* mutant and in cells expressing wild type *BEM1* treated with 1-NAA (Supplementary Fig. 3b). Thus, the SPB duplication cycle continues regardless of the severity of polarity defects, raising the question of how subsequent spindle dynamics would be affected. To address this, the time from SPB separation to anaphase onset was extracted in *BEM1* control cells and the three categories of mutant cells. In *BEM1* control cells this was 43 minutes, increasing to 64 minutes in *bem1 cdc24^lip^* mutant cells that displayed a stable pole, and to 82 minutes in *bem1 cdc24^lip^*mutants that displayed an unstable pole but eventually budded, and 177 minutes in mutant cells with an unstable pole that did not form a bud (Fig. 2b, c). Thus, the cell cycle delay correlated with the severity of the polarity defect incurred. When the *cdc28^cdk^*^1^ *Y19F* mutation was introduced into the *bem1 cdc24^lip^* mutant, the cell cycle delay was ablated, regardless of whether cells budded or not, confirming that the cell cycle delay was Swe1^Wee1^-dependent (Fig. 2b, c). Moreover, the overall percentage of multinucleate cells increased from 56% in the *bem1 cdc24^lip^* mutant to 81% in *bem1 cdc24^lip^ cdc28^cdk^*^1^ *Y19F* mutant cells (Fig. 2d). Remarkably, if the *bem1 cdc24^lip^cdc28^cdk^*^1^ *Y19F* cells displayed any instability in polarity establishment, they invariably failed to bud and all of these cells became multinucleate (Fig. 2e). These observations underscore the protective role of Cdc28^Cdk1^ inhibitory phosphorylation when polarity defects are encountered (Fig. 2e). As an additional control, deletion of the *SWE1^WEE^*^1^ gene in the *bem1 cdc24^lip^*mutant resulted in the appearance of multinucleate cells faster and to a higher extent than in the *bem1 cdc24^lip^* mutant alone (Supplementary Fig. 3c). Collectively, these results demonstrate that polarity defects are sensed and translated into a Swe1^Wee1^-dependent cell cycle delay. The extent of the temporal cell cycle delay is proportional to the severity of the polarity problem encountered.

### The extended window of G1 cyclin expression is a direct response to polarity perturbations

While a Swe1^Wee1^–dependent cell cycle delay provides a temporal window in which polarity defects can be repaired, it is unknown if Cdc28^Cdk1^ activity plays a role in the repair process itself. The observation that the G1 cyclin Cln2 was expressed during a longer temporal window in the *bem1 cdc24^lip^*mutant is consistent with a model in which G1 cyclin Cdc28^Cdk1^ activity could contribute to polarity correction, since Cdc42 module components involved in GTPase activation are specific substrates of G1 cyclin Cdc28^Cdk1^ both *in vitro* and *in vivo* ^11,16^. However, the extended duration of Cln2 expression might alternatively be an indirect consequence of Swe1^Wee1^-dependent inhibition of M-phase Cdc28^Cdk1^ activity, since the M-phase cyclin Clb2 represses G1 cyclin expression during mitosis by inhibiting the SBF/MBF activity that drives G1 cyclin transcription ^55^. Thus, when Swe1^Wee1^ is activated in the *bem1 cdc24^lip^* mutant, the repression of Cln2 transcription might be alleviated, leading to Cln2 expression, as had been suggested previously ^34^. The latter model predicts that the extended temporal window of Cln2 expression in the *bem1 cdc24^lip^* mutant would depend on Swe1^Wee1^ activity. This was tested by comparing Cln2 expression in *BEM1* control cells, *bem1 cdc24^lip^* and *bem1 cdc24^lip^ cdc28^cdk^*^1^ *Y19F* cells in which the Swe1^Wee1^ checkpoint is abrogated. Using the AID system, both Cln2 phosphorylation and its window of expression were similar in the *bem1 cdc24^lip^* mutant and in the *bem1 cdc24^lip^ cdc28^cdk^*^1^ *Y19F* double mutant (Fig. 3a and Supplementary Fig. 4a and b). Moreover, the double mutant cells became multinucleate faster and to a greater extent than the *bem1 cdc24^lip^*mutant alone within the single cell cycle of the experiment, indicative of the protective effect conferred by Swe1^Wee1^ activity (Fig. 3b). We conclude that Cln2 phosphorylation and its extended expression pattern are not indirect consequences of Swe1^Wee1^ checkpoint activity.

**Fig. 3.**
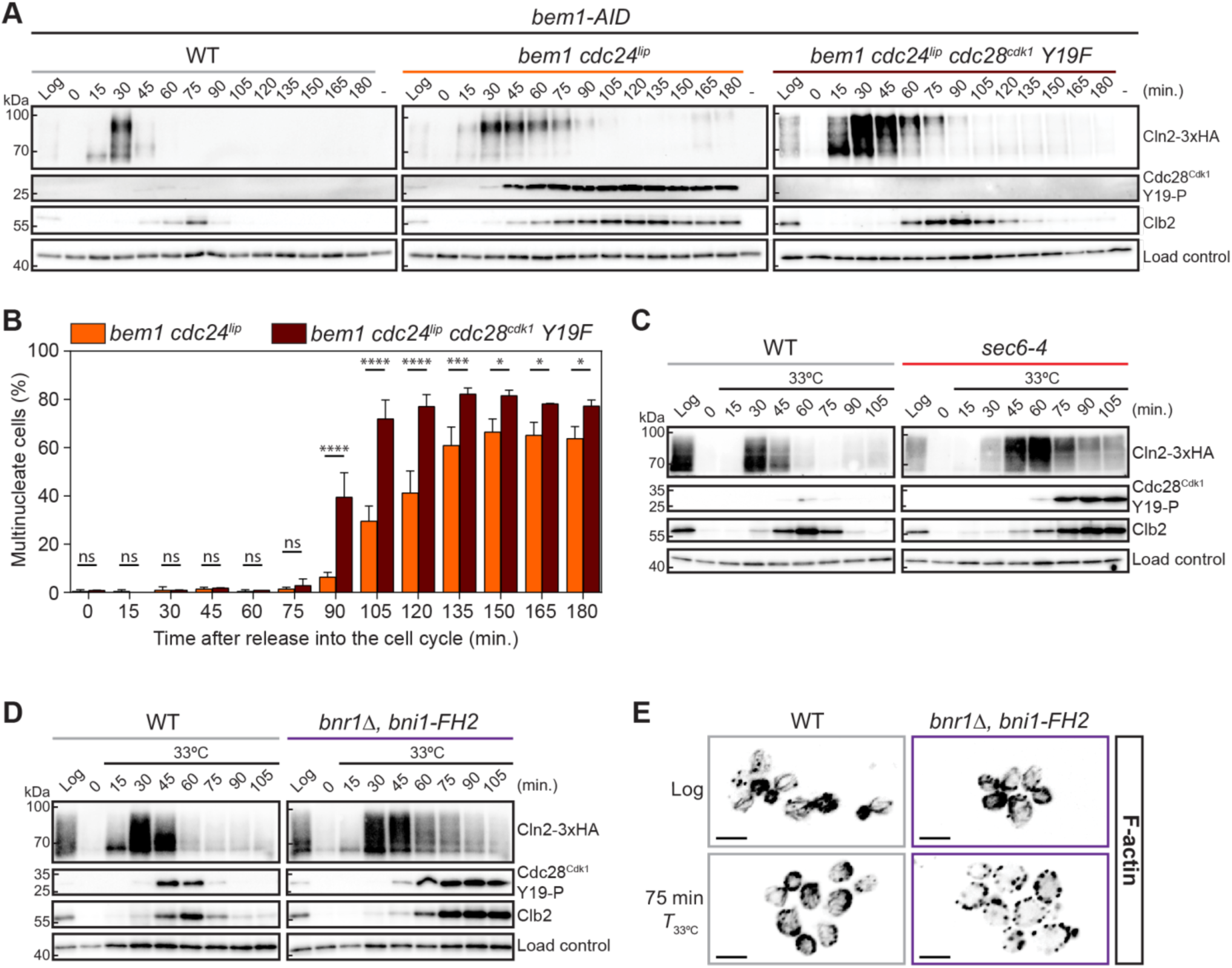
G1 cyclin hyperphosphorylation and stabilization in the *bem1 cdc24^lip^* mutant are not indirect effects of Swe1^Wee1^ activation and are evident in other polarity mutants. **(a)** Cell cycle time-course of WT, *bem1 cdc24^lip^*and *bem1 cdc24^lip^ cdc28^cdk^Y19F* cells arrested in G1 with alpha-factor, treated with auxin for the final 30 minutes of the arrest, then released into the cell cycle. Samples were removed at the indicated times. **(b)** The frequency of multinucleate cells was scored after DAPI staining (*n* > 100 cells per time point). Values display mean +/- SD. Multiple *t*-tests were performed (\**P < 0.05*, \*\*\**P < 0.001*, \*\*\*\**P* < 0.0001). **(c)** Cell cycle time-course of WT and *sec6-4* cells after release into the cell cycle at the restrictive temperature (33°C). **(d)** Cell cycle time-course of WT and *bnr1Δ, bni1-FH2* cells after release into the cell cycle at the restrictive temperature (33°C). **(e)** Maximum-intensity projected z-stacks of WT and *bnr1Δ, bni1-FH2* cells from the time-courses shown in panel d, where F-actin is visualized by Alexa-543 phalloidin staining. Fluorescence has been inverted. Scale bars = 5 µm.

This left the alternative model to test: that G1 cyclin phosphorylation and extended expression may be a direct response to polarity defects that contributes to polarity correction. Reasoning that this may be a general response to defects in polarized growth, the dynamics of Cln2 expression were monitored in the *sec6-4*, an exocyst mutant that blocks the tethering of post-Golgi vesicles to the plasma membrane when shifted to the restrictive temperature, thereby preventing bud growth. Wild type control and *sec6-4* cells were synchronized with alpha-factor and released into the cell cycle then shifted to 33°C after 15 minutes. In control cells, Cln2 expression peaked 30 minutes after release into the cell cycle, dropped at 45 minutes and was largely absent at 60 minutes, whereas in *sec6-4* cells it peaked at 60 minutes and was still detected at 75 minutes. Moreover, Cln2 displayed a slower electrophoretic mobility in the *sec6-4* mutant than in control cells, reminiscent of its hyperphosphorylation observed in the *bem1 cdc24^lip^* mutant (Fig. 3c). *sec6-4* mutant also displayed elevated Cdc28^Cdk1^ Y19 phosphorylation in comparison to control cells, indicative of Swe1^Wee1^ activation. Importantly, this Y19 phosphorylation accumulated after the appearance of phosphorylated Cln2, indicating that it is a subsequent event. The formin mutant *bnr1Δ bni1-FH2* was a third polarity mutant in which Cln2 was phosphorylated and expressed for longer than control cells after shifting cells to the restrictive temperature (Fig. 3d)^56^. In this mutant, the actin cables used for post-Golgi vesicle transport to the plasma membrane are attenuated at the restrictive temperature, thus blocking polarized growth (Fig. 3e). In conclusion, the G1 cyclin phosphorylation and extended pattern of expression observed in the *bem1 cdc24^lip^* mutant is an adaptive response to defects in polarized growth.

### G1 cyclin signaling controls Swe1^Wee1^ activity and corrects polarity errors

Since Cln2 hyperphosphorylation was the earliest cell cycle event in polarity mutants to diverge from control cells, we sought to test how it contributed to the adaptive response when polarity defects were experienced. Cln2 contains seven Cdc28^Cdk1^ consensus sites that upon phosphorylation generate a phospho-degron that is recognized by the SCF^Grr1^ E3 ubiquitin ligase, confining Cln2 expression to G1. Mutation of all seven sites stabilizes the protein and the resulting mutant localises predominantly to the nucleus ^57^. However, Cln2 is also phosphorylated extensively on sites other than Cdc28^Cdk1^ consensus sites, raising the possibility that these modifications may play other regulatory functions ^58–60^. Moreover, the phosphorylation that we observe in the *bem1 cdc24^lip^*mutant is associated with an increased window of Cln2 expression rather than its destabilization. We therefore first tested if Cln2 was stabilized in the mutant by expressing the protein from the *MET25* promoter, a conditional promoter that can be shut-off by addition of methionine (Supplementary Fig. 5a-c). Quantification of the signal from western blots indicated that the half-life of WT Cln2 was 22 minutes and 34 minutes in the *bem1 cdc24^lip^* mutant, confirming that the protein was stabilized. As an additional control for the sensitivity of the assay, we confirmed that a *cln2* mutant in which all Cdk1^Cdc28^ consensus sites were mutated (*cln2*^4T3S^) was stabilized, displaying a half-life of 56 minutes compared to 22 minutes for WT Cln2 (Supplementary Fig. 5d, e). To next investigate the role of Cln2 phosphorylation, two mutant versions of Cln2 were generated; one in which 24 phosphorylation sites mapped by mass spectrometry by us and others were mutated to alanine to block Cln2 phosphorylation (*cln2-Ala*) ^59–61^. Conversely, the same sites were mutated to aspartate to impart a negative charge resembling phosphorylation (*cln2-Asp*). The mapped phosphorylation sites cluster in regions of Cln2 that are predicted to be unstructured, at some distance from Cdc28^Cdk1^- and substrate-docking sites (Supplementary Fig. 6a-c). Moreover, only two of the seven Cdk1^Cdc28^ consensus sites that collectively form the phospho-degron for the SCF^Grr1^ complex were identified by mass spectrometry, so only these two consensus sites were mutated.

In order to test the effect of *cln2* phosphorylation on cell cycle progression and Swe1^Wee1^ activity, we transformed the G1 cyclin constructs *CLN2-3xHA, cln2-Ala-3xHA* and *cln2-Asp-3xHA* into *BEM1* cells or *bem1 cdc24^lip^* mutant cells in the AID system. Since the G1 cyclins Cln1, 2 and 3 play an essential but overlapping role in G1 progression, we also deleted *CLN1* and *CLN3* to avoid confounding effects deriving from redundancy. Cells were synchronized with alpha-factor and 30 minutes before release 1-NAA was added to degrade bem1-AID. Control experiments indicated that Bem1 was expressed comparably in the strains and that bem1-AID was degraded after 1-NAA addition (Supplementary Fig. 6d-g). In *BEM1 CLN2-3xHA* cells, successive distinct peaks of Cln2 then Clb2 expression were observed at 30 and 75 minutes after release into the cell cycle. We also observed that the Cdc42 GAP Rga2, a Cln2 substrate, became phosphorylated at 30 minutes, as Cln2 levels rose, displaying >50% dephosphorylation at 90 minutes (Fig. 4a). In *bem1 cdc24^lip^ CLN2-3xHA* cells, Cln2 was immediately phosphorylated, displaying a longer temporal window of expression. The phosphorylated Cln2 was active, since Rga2 remained phosphorylated for longer than in the *BEM1* control, peaking at 90 minutes and displaying >50% dephosphorylation at 135 minutes (Fig. 4b). As expected, the *bem1 cdc24^lip^* mutant triggered Swe1^Wee1^ activity, evident by elevated Cdc28^Cdk1^ Y19 phosphorylation. The *bem1 cdc24^lip^ cln2-Ala-3xHA* mutant strongly perturbed, but did not eliminate, cln2 phosphorylation. Rga2 phosphorylation was slightly advanced compared to *bem1 cdc24^lip^ CLN2-3xHA* cells, peaking at 75 minutes and displaying >50% dephosphorylation at 120 minutes (Fig. 4c). Levels of the M-phase cyclin Clb2 were also advanced, peaking at 90-105 minutes in *bem1 cdc24^lip^ cln2-Ala-3xHA* cells compared to 135 minutes in *bem1 cdc24^lip^ CLN2-3xHA* cells, indicating that perturbation of Cln2 phosphorylation advanced the onset of mitosis. Reduced Cdc28^Cdk1^ Y19 phosphorylation suggested mitosis was advanced in *cln2-Ala-3xHA* cells due to weakened Swe1^Wee1^ activity. Conversely, Swe1^Wee1^ activity was strongly elevated in *bem1 cdc24^lip^ cln2-Asp-3xHA* cells where Cln2 phosphorylation was mimicked. Cdc28^Cdk1^ Y19 was extensively phosphorylated and Clb2 levels did not decline during the course of the experiment, indicative of a robust mitotic delay (Fig. 4d). Quantification of Cdc28^Cdk1^ Y19 phosphorylation in the *bem1 cdc24^lip^* mutant in independent time-courses indicated that Swe1^Wee1^ activity was lowest in *BEM1* cells, increased in *cln2-Ala,* then *CLN2* cells and highest in *cln2-Asp* cells (Fig. 4e). Elevated Swe1^Wee1^ kinase activity in *bem1 cdc24^lip^ cln2-Asp* cells followed a prolonged window of cln2-Asp-3xHA expression in the mutant, resembling an exaggerated version of Cln2 expression in the *bem1 cdc24^lip^* mutant (Fig. 4d). Moreover, Rga2 exhibited prolonged phosphorylation in *bem1 cdc24^lip^ cln2-Asp-3xHA* cells, displaying >50% dephosphorylation only after 210 minutes. The robust phosphorylation of a Cln2 substrate *in* vivo indicates that like phosphorylated Cln2, the negative charge imparted by aspartate residues does not non-specifically impair Cln2 function. Since phosphorylation of the GAP Rga2 inactivates it, the results are also indicative of *bem1 cdc24^lip^ cln2-Asp-3xHA* cells mounting a prolonged self-correcting response that inactivates Cdc42 GAP activity in an attempt to generate sufficient Cdc42-GTP to support a stable polarity axis.

**Fig. 4.**
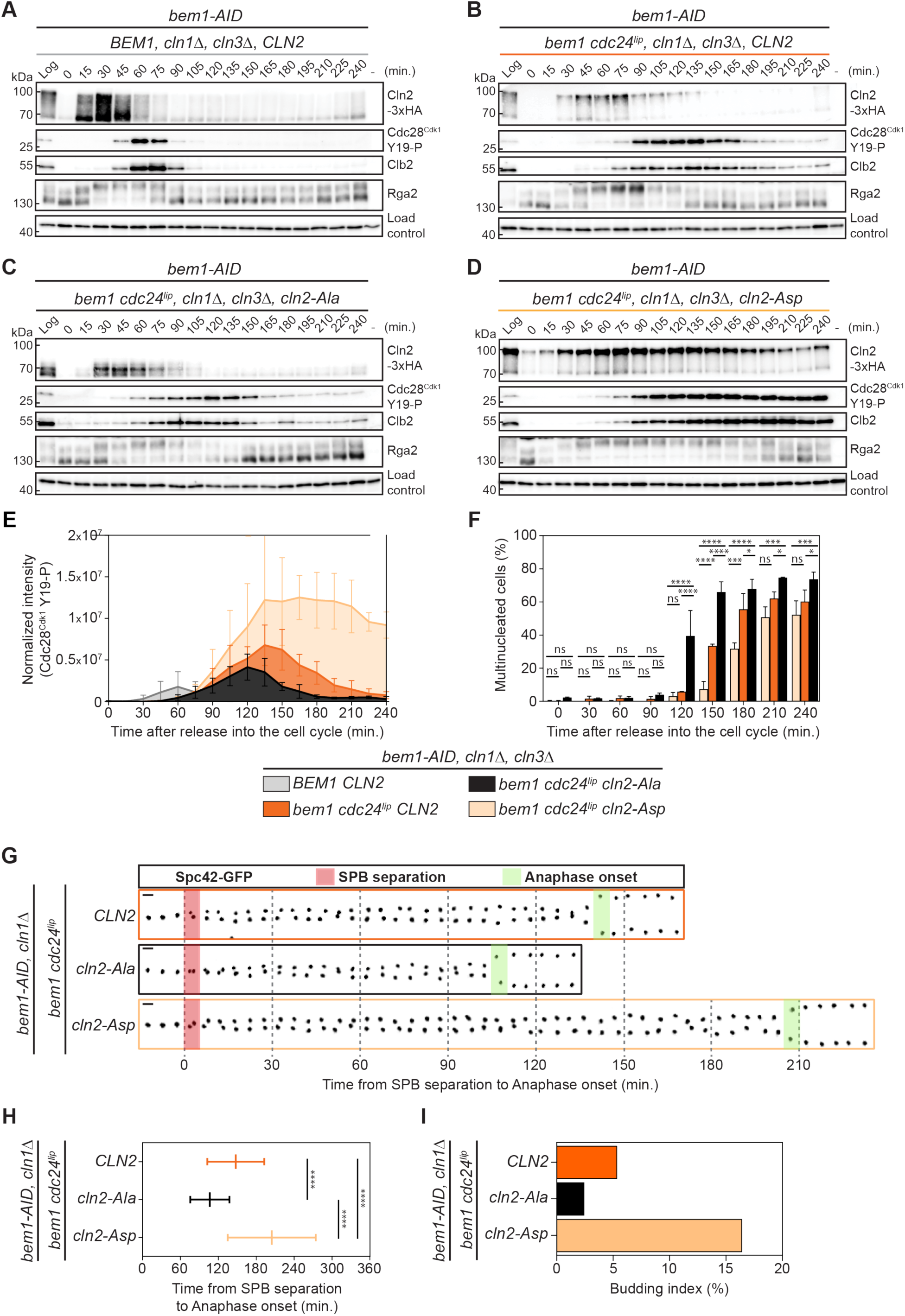
G1 cyclin signaling contributes to Swe1^Wee1^ activity to protect cells from inappropriate nuclear division. **(a)** Cell cycle time-course of control *BEM1* cells after arrest in G1, treatment with auxin for the final 30 minutes of synchronization, then release into the cell cycle. **(b)** Analysis of *bem1 cdc24^lip^ CLN2* cells. **(c)** Analysis of *bem1 cdc24^lip^ cln2-Ala* cells. **(d)** Analysis of *bem1 cdc24^lip^ cln2-Asp* cells. **(e)** Normalized intensity of Cdc28 Y19 phosphorylation signal from 3 cell cycle time-courses. **(f)** The frequency of multinucleate cells from the time-courses displayed in a-d (*n* > 100 cells per time-point). A two-way ANOVA was performed (\**P < 0.05,* \*\*\**P < 0.001*, \*\*\*\**P* < 0.0001). **(g)** Spc42-GFP dynamics in G1-arrested, auxin-treated cells. Maximum-intensity projected z-stacks of inverted fluorescence images. SPB separation is time 0 (pink) and anaphase onset is in green. Scale bars = 2 µm. **(h)** Time from SPB separation to anaphase from movies in (g) (*n* = 56 cells for *CLN2*; *n* = 82 cells for *cln2-Ala*; *n* = 67 cells for *cln2-Asp*). Values display mean +/- SD. Unpaired *t-*tests with Welch’s correction were performed (\*\*\*\**P* < 0.0001). **(i)** Frequency of budded cells from (g) (*n* = 56 cells for *CLN2*; *n* = 82 cells for *cln2-Ala*; *n* = 67 cells for *cln2-Asp*).

Concomitantly, G1 cyclin phosphorylation triggered by polarity defects feeds into the strength of subsequent Swe1^Wee1^ activity: attenuating G1 cyclin phosphorylation weakened Swe1^Wee1^ activity, while mimicking G1 cyclin phosphorylation robustly strengthened it (Fig. 4e). We therefore assessed how G1 cyclin phosphorylation and the level of subsequent Swe1^Wee1^ activity would protect cells against inappropriate nuclear division. To do so, the frequency of multinucleate cells over time was quantified as a biological readout of Swe1^Wee1^ activity. In the *bem1 cdc24^lip^ cln2-Ala* mutant, the appearance of multinucleate cells occurred faster and in greater numbers over time, consistent with weakened Swe1^Wee1^ activity (Fig. 4f). Conversely, in the *bem1 cdc24^lip^ cln2-Asp* cells, multinucleate cells appeared at a slower rate, consistent with heightened Swe1^Wee1^ activity. Kymographs of SPB dynamics using Spc42-GFP as a marker confirmed that the time from SPB separation to anaphase onset was reduced in *bem1 cdc24^lip^ cln2-Ala* cells compared to *CLN2* cells, while being delayed in the *bem1 cdc24^lip^ cln2-Asp* mutant (Fig. 4g, h and Supplementary Movie 7). We conclude that phosphorylation of the G1 cyclin Cln2 that occurs extensively in the *bem1 cdc24^lip^* mutant is involved in relaying signals to Swe1^Wee1^ that slow nuclear division, providing a temporal window for polarity correction. G1 cyclin activity also contributes directly to error correction by phosphorylating GTPase components such as the GAP Rga2. This was borne out quantitatively by comparing the percentage of budded *bem1 cdc24^lip^* mutant cells in the G1 cyclin phospho-mutants in the first cell cycle after auxin addition (Fig. 4i). Of these mutants, the *bem1 cdc24^lip^ cln2-Asp* cells that displayed the longest window of Rga2 phosphorylation, and therefore the longest period of GAP inactivation, displayed more budded cells (16%) compared with *bem1 cdc24^lip^ cln2-Ala* cells (3%).

### Polarity defects trigger the spatial reconfiguration of G1 cyclin activity

The Cdc28^Cdk1^ substrates that regulate polarity such as Rga2 are localized in the cytoplasm, whereas Cln2 localises to both the nucleus and cytoplasm. Since the nucleocytoplasmic partitioning of Cln2 has been linked to its phosphorylation, we next monitored Cln2 dynamics during the adaptive response to polarity defects ^57, 62^. This was not previously possible due to the slow maturation kinetics of GFP relative to the rapid-proteolysis of and low expression levels of Cln2. To overcome these challenges, Cln2 was tagged at its endogenous locus with sfGFP in wild type control and *bem1 cdc24^lip^* mutant cells and imaged by time-lapse spinning-disk confocal microscopy. The nucleoplasmic protein Pus1 was also tagged as a nuclear marker. The dynamics of Cln2-sfGFP were monitored in individual cells after synchronous release into the cell cycle. Cln2-sfGFP expression in control cells was evident in the cytoplasm and prominently in the nucleus for around 45 minutes, consistent with the temporal expression observed in cell populations by western blotting (Fig. 5a and Supplementary Movie 8). In contrast, nuclear enrichment of Cln2-sfGFP expression was not observed in *bem1 cdc24^lip^* mutant cells, which instead displayed a prominent and sustained burst of cytoplasmic Cln2. In a *bem1 cdc24^lip^* mutant cell that formed a bud, Cln2-sfGFP was rapidly degraded after bud emergence, indicating that successful polarization and budding is linked to Cln2 degradation (Fig. 5a middle panels and Supplementary Movie 9). However, the majority of *bem1 cdc24^lip^* mutant cells did not form a bud and these cells displayed a continuous increase in cytoplasmic Cln2-sfGFP expression that eventually declined (Fig. 5b and Supplementary Movie 10). Fluorescence intensity quantification indicated that *bem1 cdc24^lip^* mutant cells expressed higher global levels of Cln2-sfGFP at steady-state, as well as displaying higher fluorescence intensity in the cytoplasm and a lower nuclear:cytoplasmic ratio than control *BEM1* cells (Supplementary Fig. 7a-c). This spatial reconfiguration of G1 cyclin activity from the nucleus to the cytoplasm in response to polarity defects was also evident by monitoring the dynamics of the *cln2* phosphorylation mutants. When polarity defects were induced in *bem1 cdc24^lip^* cells, *cln2-Ala-sfGFP* mutants exhibited more prominent nuclear fluorescence than in *bem1 cdc24^lip^*cells alone, consistent with the non-phosphorylated form of the protein being enriched in the nucleus (Fig. 5c and Supplementary Movie 11). Conversely, in *bem1 cdc24^lip^* cells, *cln2-Asp-sfGFP* displayed an increased, sustained burst of cytoplasmic fluorescence and partial exclusion from the nucleus that was evident from quantification (Fig. 5c-f and Supplementary Movie 12). These results are consistent with previous reports that dephosphorylated Cln2 was enriched in nucleus of early G1 cells while phosphorylation was associated with its cytoplasmic localization ^57,62^. Moreover, the cytoplasmic enrichment of Cln2 was observed in other mutants defective in polarized growth such as *sec6-*4 (Supplementary Fig. 7d-h). The data are indicative of the re-routing of Cln2 to the cytoplasm where it accumulates, activating Cdc42 as an adaptive response in the *bem1 cdc24^lip^* mutant and other polarity mutants.

**Fig. 5.**
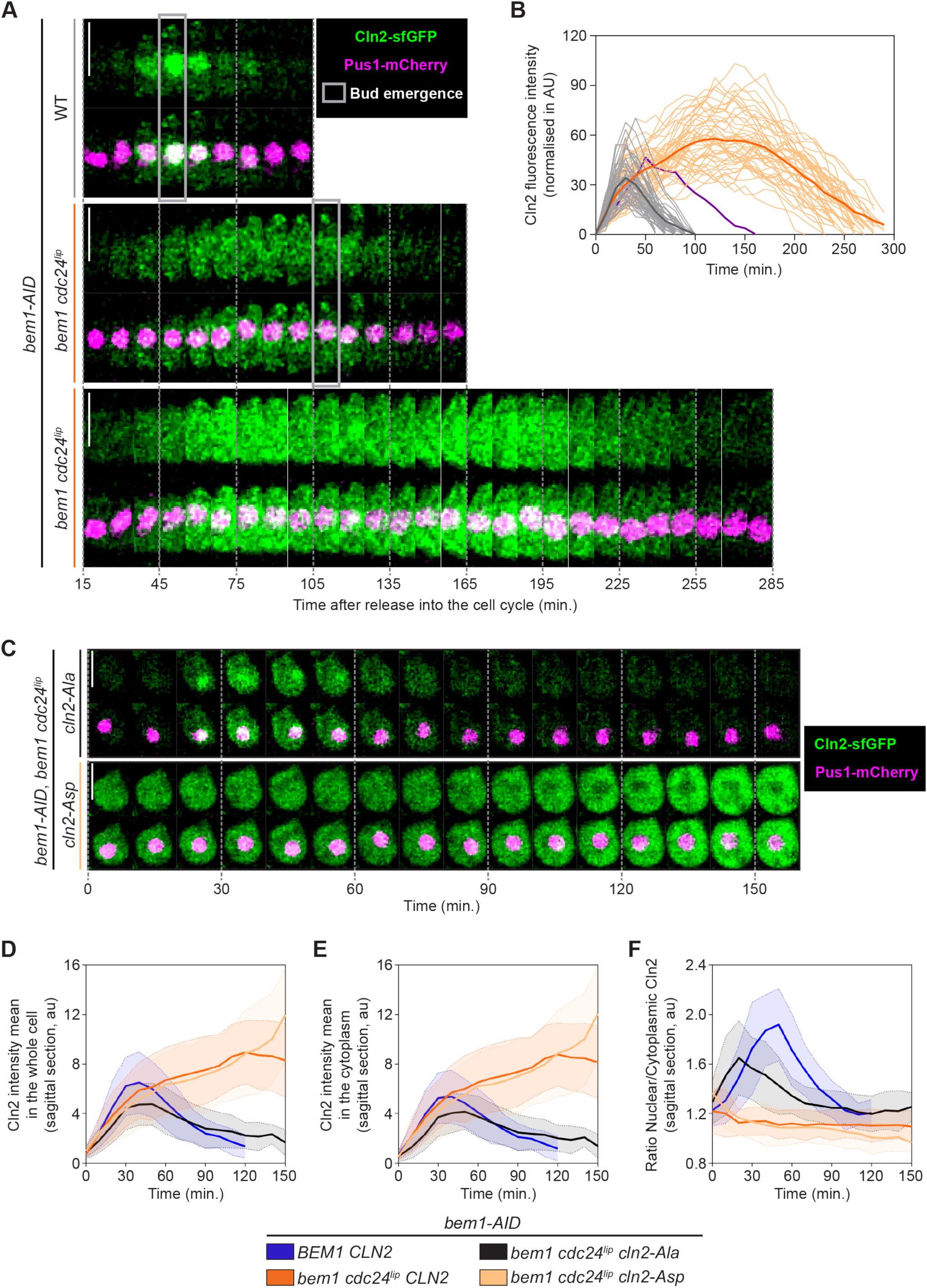
The *bem1 cdc24^lip^* mutant triggers an adaptive response that augments cytoplasmic G1 cyclin levels. **(a)** *BEM1* and *bem1 cdc24^lip^* mutant cells expressing Cln2-sfGFP and Pus1-mCherry (nucleoplasm) were arrested in G1 with alpha-factor, treated with auxin for the final 30 minutes of the arrest, then released into the cell cycle. Cln2-sfGFP dynamics were visualized at 10-minute intervals. Images are maximum-intensity projected z-stacks of Cln2-sfGFP (green) and Pus1-mCherry (magenta) where bud emergence (grey rectangle) is indicated. Time 0 indicates release into the cell cycle. Scale bars = 5 µm. An example of *bem1 cdc24^lip^* mutant cells that buds (middle) and that does not bud (bottom) is shown. **(b)** Quantification of Cln2 fluorescence intensity in each time frame in WT cells (*n* = 47) and *bem1 cdc24^lip^* mutant cells (*n* = 48). Each line is an individual cell, and bold lines display the mean of WT and *bem1 cdc24^lip^* mutant cells. The purple line displays Cln2 dynamics in a *bem1 cdc24^lip^* mutant cell that buds, shown in panel (a, middle). **(c)** *bem1 cdc24^lip^ cln2-Ala* (upper panel) and *bem1 cdc24^lip^ cln2-Asp* (lower panel) were arrested in G1 with alpha-factor, treated with auxin for the final 30 minutes of the arrest, then released into the cell cycle. Cln2-sfGFP dynamics were visualized at 10-minute intervals. Images are maximum-intensity projected z-stacks of Cln2-sfGFP (green) and Pus1-mCherry (magenta). Scale bars = 5 µm. **(d)** Quantification of Cln2 mean fluorescence intensity in the whole cell in WT cells (*n* = 47), *bem1 cdc24^lip^* mutant cells (*n* = 47), *bem1 cdc24^lip^ cln2-Ala* mutant cells (*n* = 51) and *bem1 cdc24^lip^ cln2-Asp* mutant cells (*n* = 50). Values display mean +/- SD. **(e)** Quantification of Cln2 mean fluorescence intensity in the cytoplasm in WT cells (*n* = 47), *bem1 cdc24^lip^* mutant cells (*n* = 47), *bem1 cdc24^lip^ cln2-Ala* mutant cells (*n* = 51) and *bem1 cdc24^lip^ cln2-Asp* mutant cells (*n* = 50). Values display mean +/- SD. **(f)** The ratio of nuclear to cytoplasmic Cln2 is displayed for WT cells (*n* = 47), *bem1 cdc24^lip^* mutant cells (*n* = 47), *bem1 cdc24^lip^ cln2-Ala* mutant cells (*n* = 51) and *bem1 cdc24^lip^ cln2-Asp* mutant cells (*n* = 50). Values display mean +/- SD.

## Discussion

The nanoclustering of proteins on the plasma membrane was previously modeled to digitize signaling, contributing to the decisiveness of Ras and Rho GTPase responses ^25^. In this study, interfering with Cdc42 nanoclustering on the plasma membrane by ablating multivalent anionic lipid interactions underscored its importance for polarity dynamics *in vivo*; polarity establishment was delayed and was not decisive, but rather vacillated, displaying strong defects in maintenance. However, the loss of decisiveness ensuing from the lipid tethering defects of Cdc42 activators extended beyond GTPase signaling. Unexpectedly, the switch-like properties of the entire cell cycle control system, which is usually characterized by successive temporal waves of cyclin activity, became compromised.

The loss of the cell cycle’s switch-like characteristics derived from the activation of an adaptive response that cells mount to correct polarity defects. This involved a prolonged burst of cytoplasmic G1 cyclin expression that phosphorylated and inactivated a Cdc42 GAP, thus boosting Cdc42-GTP levels. This spatial response also fed into a temporal response that activated Swe1^Wee1^ kinase activity, delaying nuclear division via M-phase cyclin stabilization until polarity was corrected (Fig. 6). Interfering with either protective mechanism compromised euploidy. The study also illustrates how G1 cyclins not only trigger an essential cell cycle event - polarity establishment, but also monitor its successful completion. This requires G1 cyclins to be embedded within feedback networks that adjust cyclin activity and localization until the events that they monitor are executed appropriately. The strategy of upregulating the activator of a system in response to a deficit of the system’s output seems ingenious. The necessity for G1 cyclin upregulation to rectify polarity defects may derive from the specificity that G1 cyclins display for Cdc42 module components ^11,16^. However, the cost of this strategy is that G1 cyclin upregulation commits cells to one cycle of cell division. This means that even if the cell cannot repair the problem, it will nevertheless eventually undergo nuclear division, generating a multinuclear cell. This underscores the perils of passage through G1 and irreversible cell cycle commitment. It may also explain the high frequency with which G1 control is found to be aberrant in tumours ^63^. The short-term response identified in this study occurs in the cell cycle during which the polarity problem is encountered. However, in the longer-term, cells in which the scaffold *BEM1* has been deleted adopt a different strategy to mitigate the ensuing deficit of active Cdc42. These cells down-regulate inhibitors of Cdc42 due to a selective pressure for the mutation of GAPs, instead of upregulating an activator, highlighting the capacity of GTPase signaling modules to rapidly adapt in response to perturbations ^64^. The present study illustrates how these perturbations to GTPase signaling extend beyond cell polarity to affect the cell cycle control network, whose timely response is essential to coordinate cell polarity and cell cycle progression, a requirement for healthy cell proliferation.

**Fig. 6.**
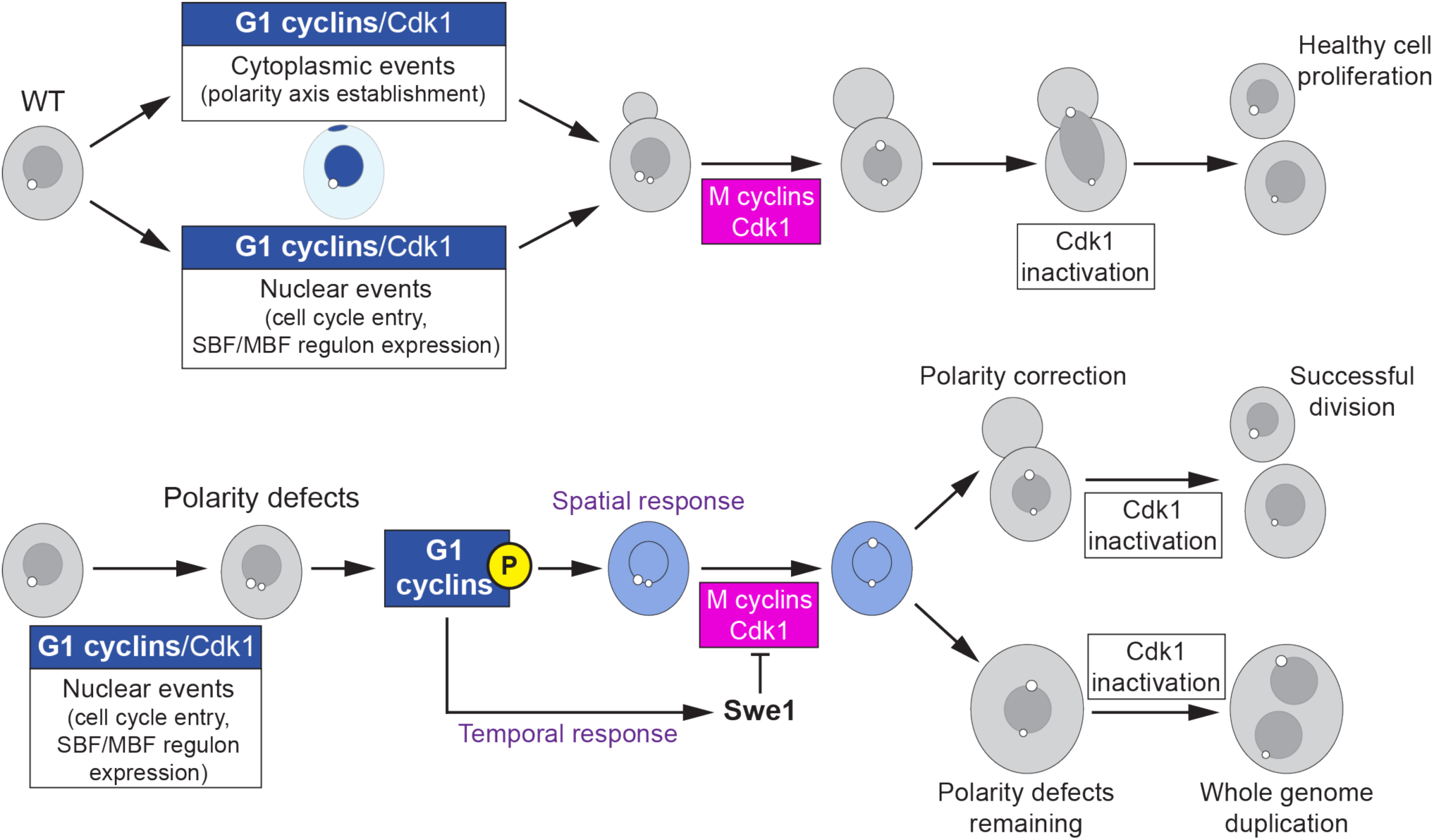
The *bem1 cdc24^lip^* mutant triggers an adaptive response that augments cytoplasmic G1 cyclin levels. Schematic model of key cell cycle events related to G1 in wild type cells (upper panel) and the corrective events that are triggered in response to polarity defects including the outcome of these events (lower panel).

## Methods

### Plasmid construction

Plasmid names are provided in table S1. To construct AID strains, pDM1052 was generated by cutting pDM443 (pRS303) with XhoI and SacI and ligating *pBEM1-bem1 bc-14E + px (K338M, K348A, R349A & R369A)-Pk3*. pDM1053 was generated by ligating *pBEM1-BEM1-Pk3* (XhoI-EcoRI) into pRS303. pDM1052 and pDM1053 were integrated at the *his3* locus by cutting with NheI. Synthesized gene fragments encoding *cln2-Ala-3xHA* (*S233A, S236A, S239A, T242A, T378A, S380A, T381A, S383A, S385A, S389A, S391A, T392A, S393A, S396A, S399A, S400A, S401A, S403A, T468A, S472A, T476A, S528A, S529A, S530A)* or *cln2-Asp-3xHA (S233D, S236D, S239D, T242D, T378D, S380D, T381D, S383D, S385D, S389D, S391D, T392D, S393D, S396D, S399D, S400D, S401D, S403D, T468D, S472D, T476D, S528D, S529D, S530D)* containing NdeI and SpeI sites were cloned into pDM64 (*pCLN2-CLN2-3xHA LEU2 (INT)*). The *cln2-3xHA* constructs were then sub-cloned into pRS406 (*URA3*) to generate pDM1085 (*pCLN2-CLN2-3xHA*), pDM1087 (*pCLN2-cln2-Ala-3xHA*) and pDM1090 (*pCLN2-cln2-Asp-3xHA*) and amplified using oligonucleotides that targeted integration at the *CLN2* locus. Integration was verified by PCR and sequencing. Gene synthesis was used to generate *cdc28^cdk^*^1^ *Y19F* mutants. *pCDC28-cdc28^cdk1^Y19F-cdc28term::natNT2* was synthesized (pDM1091), then digested with BglII and EcoRI to change the selection marker, generating pUC57-*pCDC28-cdc28^cdk1^Y19F-CDC28term::kanMX6* (pDM1092). pDM996 (pFA6a-*GAGAGA-mRuby2::HIS3)* was generated by replacing GFP in pFA6a-*GFP(S65T)::HIS* with GAGAGA-mRuby2 using PacI and AseI sites. pDM1064 and pDM1065 were generated by replacing *HIS3* in pFA6a-*GFP(S65T)::HIS3* with *CaURA3* and *natNT2*, respectively, after digestion with PmeI and BglII. pDM1140 (pFA6a-*mScarletl3-mScarletl3::HIS3*) was generated by replacing GFP in pFA6a-*GFP(S65T)::HIS3* sequentially with *mScarletl3* coding regions. All plasmids were checked by sequencing.

### Yeast strains and growth conditions

Strain names and numbers are provided in table S2. One-step PCR-based gene replacement was used for tagging at endogenous loci. To check for integration at the desired locus, flanking oligonucleotides were designed and used with untagged control DNA. Cells were grown on rich medium YPD or synthetic medium at 25°C and maintained in semi-logarithmic growth phase (OD_600 nm_ < 0.8 ml^−1^). AID strains were generated as follows: the pTIR1 plasmid was digested using PmeI, to release TIR1 for integration at the *LEU2* locus ^52^. In a second step, *BEM1* was tagged by homologous recombination (*BEM1-AID-6xFLAG::hphNT1*) at the endogenous *BEM1* locus. Finally, a WT version of *BEM1* (*his::BEM1-Pk3::HIS3*) or a mutant version of *bem1^lip^* (*his3::BEM1p-bem1 bc-14E + px (K338M, K348A, R349A & R369A)-Pk3::HIS3*) was integrated at the *his3* locus. Transformants were tested for 1-NAA sensitivity and verified by PCR and DNA sequencing.

### Testing G1 arrest of cells with alpha-factor - “halo assays”

Cells were grown at 25°C to semi-logarithmic growth phase. Two dilutions of cells were spread on YPD plates. Sterile filter paper was then applied with different concentrations of alpha-factor (0.5 mg.ml^−1^, 0.05 mg.ml^−1^ or 0.005 mg.ml^−1^). After 2-days at 25°C, halos of growth inhibition due to cell cycle arrest were observed around the filter paper (Extended Data Fig. 1d).

### Cell cycle time-courses

Cells were grown at 25°C until early-logarithmic phase, then diluted to an OD_600 nm_ of 0.3 ml^−1^. Cells were then synchronized by adding alpha-factor to 7 µg.ml^−1^ (or 0.5 µg.ml^−1^ for *bar1Δ* cells) for 3-hours at 25°C. After washing cells in YPD using a vacuum pump attached to a manifold with a 0.22 µm nitrocellulose membrane without allowing them to dry, cells were released synchronously into the cell cycle. Cells were then incubated at 25°C with agitation, and samples were taken every 15 minutes. To ensure that all cells completed a single cycle, alpha-factor (15 µg.ml^−1^) was added back to cells at 50 minutes to re-arrest them at the end of a single cell cycle. When using the AID system, bem1-AID was degraded by the addition of 0.5 mM 1-NAA, 30 minutes before washing away the alpha-factor. The same concentration of 1-NAA was added throughout the rest of the experiment to degrade any newly synthesized bem1-AID.

### Determination of protein stability

Cells expressing *MET25p-3xHA-CLN2* at the endogenous locus were grown at 25°C in YPD supplemented with 0.08 g.l^−1^ methionine until mid-logarithmic phase (Extended Data Figs. 5a-c). Cells were then washed and resuspended in minimal medium lacking methionine for 2 hours to induce expression from the MET25 promoter, before YPD + 0.08 g.l^−1^ methionine was added to repress expression. Cell lysates were analyzed by immunoblotting using anti-HA antibodies. When using the AID system, bem1-AID was degraded by the addition of 0.5 mM 1-NAA, 30 minutes before washing away the methionine. The same concentration of 1-NAA was added throughout the rest of the experiment to degrade newly synthesized bem1-AID.

“Gal-shut-off” experiments were used to determine protein stability in Extended Data Figs. 5d-e. Cells were grown in YEP medium supplemented with 2% ethanol + glycerol at 25°C until mid-logarithmic phase then diluted to an OD_600 nm_ of 0.4 ml^−1^. Galactose was then added to a final concentration of 2% to induce expression from the GAL1 promoter. After 60 minutes of gal-induction, cells were collected by centrifugation and resuspended in YEP with 2% dextrose to repress expression.

### Western blotting

Samples for Western blotting were prepared by collecting 1 OD_600 nm_ of mid-logarithmic phase cells, adding glass beads and flash freezing in liquid nitrogen. Some specific proteins required a different volume of cells to be frozen, which varied according to the sensitivity of the antibody being used for detection and protein abundance (0.5 OD_600 nm_ units for cyclins, 1.5 OD_600 nm_ units to detect phosphorylation on Y19 of Cdc28, or 2 OD_600 nm_ units for Rga2). Samples were vigorously agitated in 100 µL SDS sample buffer (65 mM Tris pH 6.8, 3% SDS, 10% glycerol, 5% β-mercaptoethanol, 100 mM β-glycerolphosphate, 50 mM NaF) supplemented with 1 mM fresh PMSF. Samples were immediately boiled and analyzed by SDS–PAGE, Western blotting, and probed with appropriate antibodies. The primary antibodies used were anti-HA (12CA5), anti-TAP (IgG peroxidase, Jackson) anti-FLAG (M2), anti-c-MYC (9E10), anti-V5 (Pk3), anti-Clb2, anti-Swe1, anti-Gin4, anti-Ade13, anti-Rga2, anti-Y19P (phospho-cdc2 (Y15) Cell Signaling Technology). In figures, logarithmically-growing samples prior to synchronization are labelled “Log”, whereas negative controls (gene deletions or untagged controls) are labelled with a minus sign (“-”).

Where anti-Cdc28^Cdk1^ phospho-Y19 chemiluminescence signals were quantified, the same membrane was used for quantification and for the measurement of a loading control (anti-Ade13). Non-specific background was removed globally using ImageLab software, and volume tools were used to measure adjusted signal intensity before normalizing anti-Cdc28^Cdk1^ phospho-Y19 to the anti-Ade13 loading control.

### Cell imaging

Cells to be imaged were grown at 25°C in synthetic medium to semi-logarithmic phase and treated, or not, with alpha-factor and 1-NAA as indicated in figure legends. Cells were mounted on 0.5% agar pads and sealed with halo-carbon oil for imaging. Live-cell imaging was carried out in a temperature-controlled room (22-25°C) using a Ti-DH inverted microscope (Nikon) equipped with a ×100 oil Plan-Apochromat objective lens (N/A 1.45), a spinning disk (CSU-W1-T1, Yokogawa), and a sCMOS camera (Roper Scientific). The system was controlled by Metamorph software (Molecular Device). Depending on the number of acquisitions, total acquisition time and cell size, between 10 and 15 Z-stacks with an interval of 0.3 or 0.5 µm were acquired.

Where necessary, cells were fixed with 3.7% formaldehyde for 20 min. After washing in PBS three times, cells were incubated for one hour with 10 µL of DAPI solution (Invitrogen) to measure the percentage of multinucleate cells (Figs. 3b and 4f). To visualize actin filaments, fixed cells were labelled with Alexa 546–phalloidin before imaging in PBS (Fig. 3e).

### Image analysis and data processing

Fiji (National Institutes of Health) was used for image analysis and fluorescence quantification. The background level of the camera was subtracted before quantification. For qualitative analyses, the timing of cell cycle events in mutant strains were compared with control cells. To better visualize these events, the images shown in Figs. 1a, 1d, 1h, 2b, 4g, 5a, 5c and Extended Data Figs. 7d, 7e were median filtered (filter size 1 × 1), whereas quantitative analysis was performed on raw data.

Sec3-GFP polarization was defined when the signal accumulated at the plasma membrane for at least 10 minutes (Figs. 1, 2 and Extended Data Figs. 1, 3). Cell separation and bud emergence were defined from DIC images. The duration of Clb2 and Clb5 expressions was defined qualitatively when a signal stronger than background was observed (Fig. 1 and Extended Data Fig. 1). SPB separation was defined when the 2 SPBs were resolved, and anaphase onset was defined when the distance between the two SPBs doubled in one frame (Figs. 2 and 4).

In Fig. 5b, Cln2 fluorescence intensity was measured in the whole cell on sum-intensity projected z-stacks in each time frame, where fluorescence intensity values were extracted after background subtraction and Pus1 levels, which are not cell cycle regulated, were used to normalize for bleaching. In contrast, Cln2 fluorescence intensity was measured in sagittal sections through the cell in each time frame in Figs. 5d-f, where fluorescence intensity values were extracted after background subtraction and Pus1 levels were used to normalize for bleaching as well as delineating nuclear and cytoplasmic signals. A similar strategy was used in Extended Data Fig. 7, where fluorescence intensity values were extracted after background subtraction, and nuclear and cytoplasmic signals were measured using the Pus1 signal to delineate the nucleoplasm.

### Statistical analysis

All statistical analyses were performed with Prism software. Means ± SD are shown in all graphs. In Extended Data Fig. 1e and 4b, the curves were smoothed using LOWESS analysis (medium). In Figures 1, 2 and Extended Data Fig. 7, all individual values are also represented by points. After testing for the normal distribution of data, *t*-tests with Welch correction or Mann-Whitney tests were performed (\**P* < 0.05, \*\**P* < 0.01, \*\*\**P* < 0.001, \*\*\*\**P* < 0.0001). The measurement of multinucleated cell percentages was repeated 3 times (Fig. 1i) and all the cells were grouped after ensuring that these repeat experiments were not significantly different. Two-way ANOVA in Fig. 1e, 4d, Extended Data Fig. 3c or multiple *t*-tests in Fig. 3b were performed to compare means at each time point. Half-lives of Cln2 were calculated after nonlinear regression fittings (one phase decay or sigmoidal (4PL)), and qualities of fittings are mentioned in Extended Data Fig. 5.

### Cln2 purification and mass spectrometry

3xHA-Cln2-Cdc28^Cdk1^ was purified from a yeast strain expressing the construct from the *GAL1* promoter integrated at the *CLN2* locus. The active complex was purified using rabbit polyclonal anti-HA antibody and gently eluted using HA-dipeptide and kinase assays were set up as described previously ^11^. Coomassie-stained SDS-PAGE gel slices containing Cln2 were excised, digested with trypsin and analyzed by liquid chromatography tandem mass spectrometry (LC-MS/MS) using an LTQ-FThybrid linear ion trap-Fourier transform ion cyclotron resonance (FTCR) mass spectrometer (Thermo-Electron, San Jose, CA), as described previously ^65^. Database searches were then performed against the *S. cerevisiae* Cln2 protein using the SEQUEST search algorithm and subsequently validated using an in-house software package ^66^.

## Supporting information

Peyran supplemental data

## Acknowledgments

We are grateful to Carilee Rodwell for mass spectrometry analysis of Cln2 and Cameron Mackereth for generating the Cln2 structural model. We also thank Doug Kellogg, Benoit Pinson and Bjoern Lillemeier for generously providing antibodies. We are also grateful to Simonetta Piatti, Gilles Charvin, Frank Uhlmann, Manos Mavrakis, Doug Koshland, Helle Ulrich and Isabelle Sagot for plasmids and strains. Aurélie Massoni-Laporte, Alice Fraigner, Marie-Charlotte Claverie and Noluen Touly are thanked for technical assistance. Structural models of Bem1, Cdc24 and Cln2 were generated using Alphafold. This work was supported by Agence Nationale de la Recherche ANR-21-CE13-0045 Pebbles (DMcC and AR), subventions from “La Ligue contre le cancer” (DMcC), Ministère de l’Enseignement Supérieur et de la Recherche (LP), Fondation ARC pour la recherche sur le cancer 4^th^ year PhD fellowship (LP) and Centre National de la Recherche Scientifique France (DMcC and AR).

## Author contributions

Conceptualization: DMcC. Methodology: DMcC, LP, SPG. Investigation: LP, CL, DMcC, SPG. Funding acquisition: DMcC, LP, AR. Supervision: DMcC. Writing – original draft: DMcC. Writing – review & editing: DMcC, SPG, LP, CL, AR.

## Materials & correspondence

Derek McCusker.

