## Supplementary material for "A cyclin-polarity feedback network ensures healthy cell proliferation": Peyran supplemental data

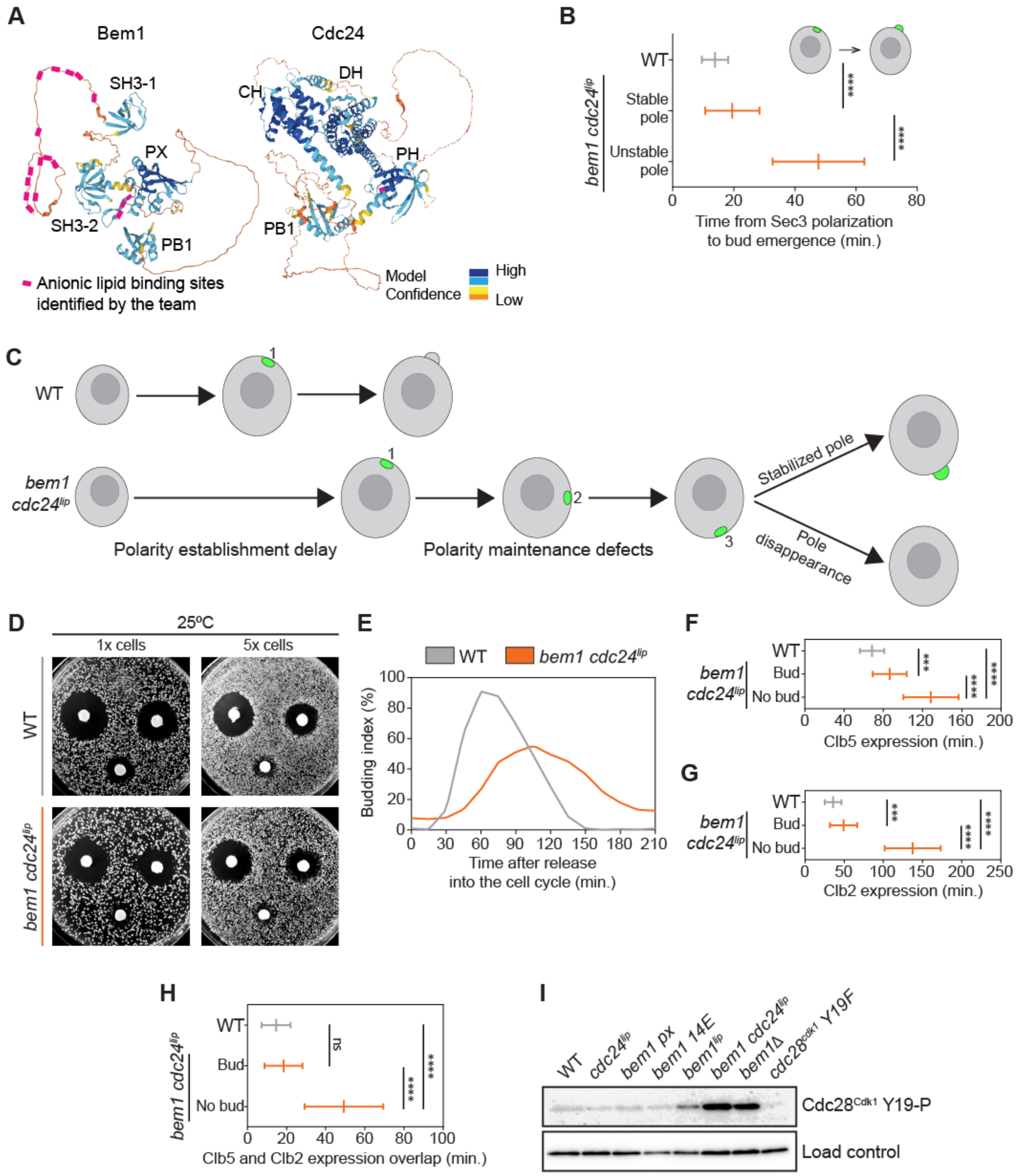


Supplementary Fig.1

**Supplementary Fig. 1. Lipid tethering defects in the *bem1 cdc24$^{lip}$* mutant delay bud emergence, impact the decisiveness of cell cycle events and trigger Swe1$^{Wee1}$ activation.**

**(a)** Alphafold models of Bem1 (https://alphafold.ebi.ac.uk/entry/P29366) and Cdc24 (https://alphafold.ebi.ac.uk/entry/P11433) in which mutated basic residues that were demonstrated to interact with anionic lipids are shown in pink. Model confidence values are included. **(b)** Time between Sec3-GFP polarization and bud emergence in wild type (*n* = 73 cells) and *bem1 cdc24$^{lip}$* cells exhibiting a stable pole (*n* = 61 cells) or an unstable pole (*n* = 18). **(c)** A schematic summarizing the polarity establishment and maintenance defects displayed by the *bem1 cdc24$^{lip}$* mutant. **(d)** Wild type and *bem1 cdc24$^{lip}$* cells were adjusted to the same $OD_{600nm}$ and grown on YPD plates containing alpha-factor at 0.5 mg.ml$^{-1}$, 0.05 mg.ml$^{-1}$ and 0.005 mg.ml$^{-1}$. Alpha-factor induces a G1 arrest that generates a halo of growth inhibition, enabling the sensitivity to G1 arrest to be assessed qualitatively after two days of growth at 25ºC. **(e)** Frequency of budded wild type and *bem1 cdc24$^{lip}$* cells at the times indicated after synchronous release into the cell cycle. This data is from the time-course shown in Figure 1F and G. **(f)** Timing of Clb5-sfGFP (S-phase) expression in wild type (*n* = 35 cells) and *bem1 cdc24$^{lip}$* cells that bud (*n* = 19 cells) and that do not bud (*n* = 27 cells). **(g)** Timing of Clb2-2xmScarletl3 (M-phase) expression in wild type (*n* = 28 cells) and *bem1 cdc24$^{lip}$* cells that bud (*n* = 18 cells) and that do not bud (*n* = 13 cells). **(h)** Timing of Clb5-sfGFP and Clb2-2xmScarletl3 expression overlap in wild type (*n* = 38 cells), budded *bem1 cdc24$^{lip}$* cells (*n* = 19 cells) and unbudded *bem1 cdc24$^{lip}$* cells (*n* = 27 cells). **(i)** Western blots of the strains indicated that were probed for Cdc28$^{Cdk1}$ Y19 phosphorylation. Values display mean +/- SD. Mann-Whitney tests were performed. (****P* < 0.001, *****P* < 0.0001).

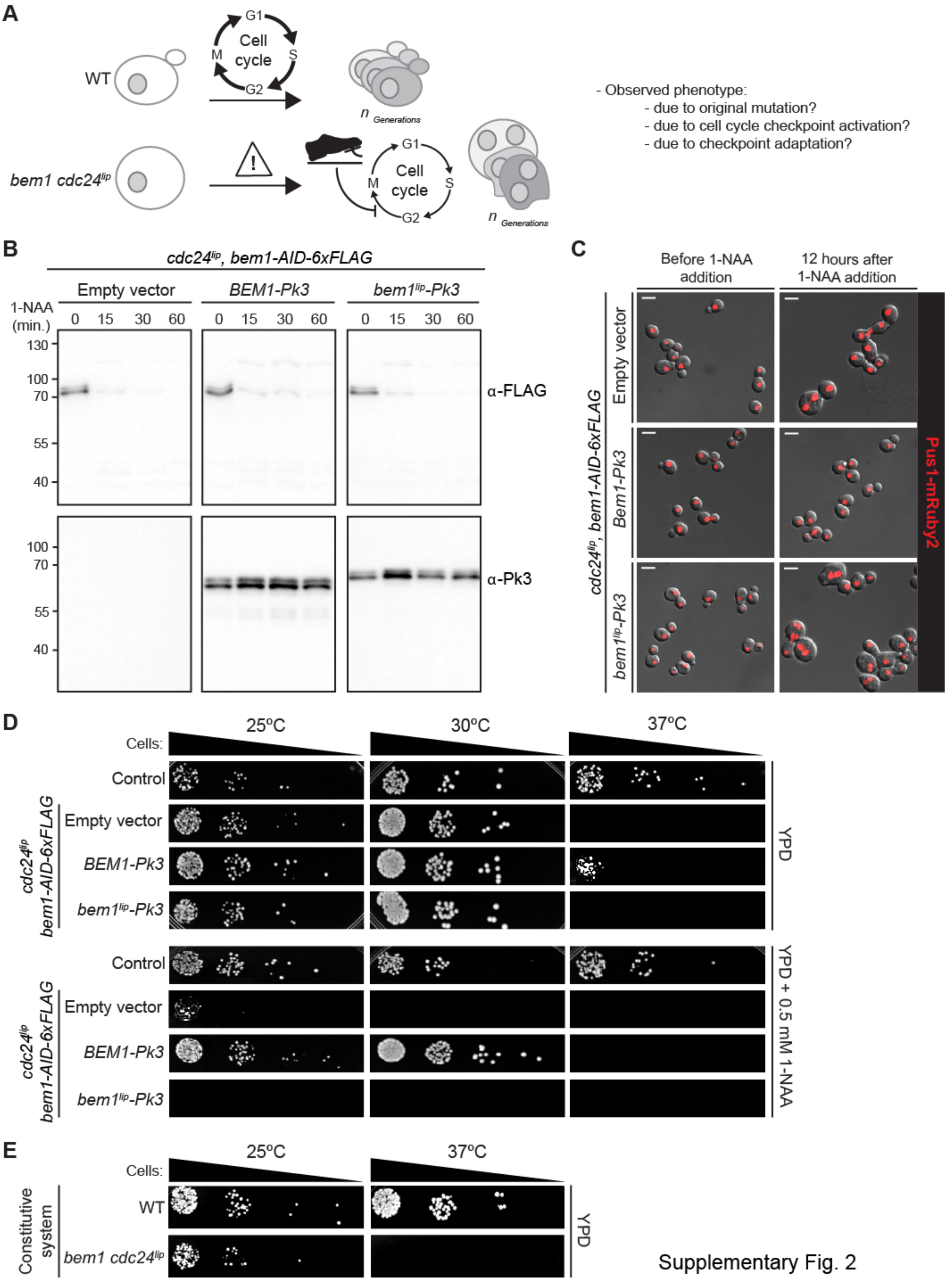


Supplementary Fig. 2

**Supplementary Fig. 2. Characterization of an auxin inducible degron (*bem1-AID*) to rapidly and conditionally ablate Bem1 function.**

**(a)** Schematic depicting the difficulty of assigning a phenotype to mutants that activate cell cycle checkpoints. **(b)** Western blots where *BEM1-Pk3*, *bem1$^{lip}$-Pk3*, or cells transformed with an empty vector were treated with the synthetic auxin hormone 1-NAA for the times indicated. Endogenous bem1-AID-6xFLAG was degraded within 15 minutes, while Bem1-Pk3 and bem1$^{lip}$-Pk3 were expressed to comparable levels, both being expressed from the *BEM1* promoter. Note that the bem1$^{lip}$-Pk3 protein displays reduced electrophoretic mobility than Bem1-Pk3 by SDS-PAGE due to the charge imparted by the replacement of 14 N-terminal lysine/arginine residues with glutamate. **(c)** Cells of the indicated genotype were treated with 0.5 mM auxin for 12 hours and Pus1-mRuby2, a marker of the nucleoplasm, was imaged. Scale bar = 5 µm. Cells containing an empty vector or *bem1 cdc24$^{lip}$* displayed morphological defects and many became multinucleate, while those transformed with wild type *BEM1-Pk3* did not. **(d)** 10-fold serial dilutions of the indicated strains were spotted onto YPD plates or YPD plates supplemented with 0.5 mM 1-NAA and grown at 25ºC, 30ºC or 37ºC for two days. When spotted onto plates in the absence of 1-NAA, all strains formed colonies at a similar rate at 25ºC and 30ºC. However, when cells containing an empty vector or the *bem1 cdc24$^{lip}$* mutant were incubated for two days in the presence of 1-NAA, they lost viability at 25ºC and 30ºC. **(e)** 10-fold serial dilutions of wild type and *bem1 cdc24$^{lip}$* cells (constitutive system) were spotted onto YPD plates and grown for two days at the temperatures indicated. We interpret the more severe loss of viability at 25ºC in the AID system (d) compared to the constitutive system (e) as being indicative of adaptive mechanisms responding to the loss of bem1 and cdc24 lipid binding in the constitutive system. The advantage afforded by the AID system is that there is insufficient time for adaptation.

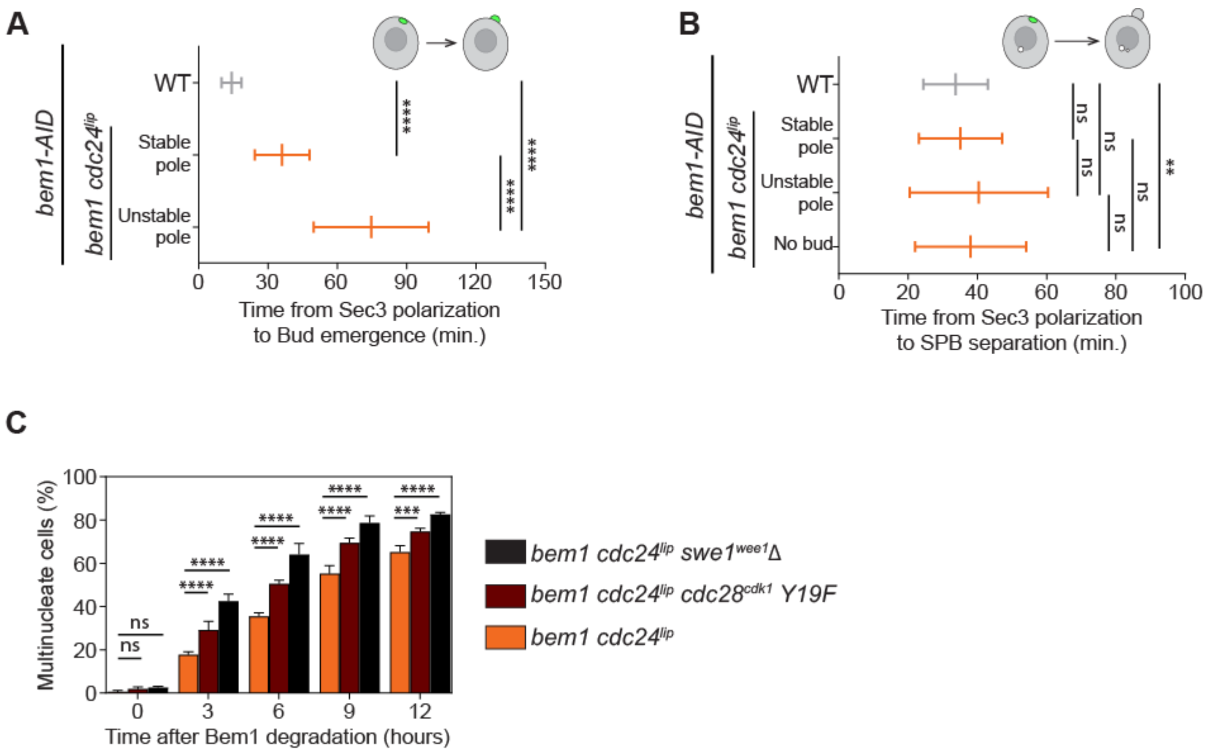


Supplementary Fig. 3

**Supplementary Fig. 3. The SPB duplication cycle continues regardless of the severity of polarity defects, resulting in the appearance of multinucleate cells.**

**(a)** Time between the first Sec3 polarization event and bud emergence in control cells (n = 190), *bem1 cdc24$^{lip}$* cells that displayed a stable pole (*n* = 43), or unstable pole (*n* = 17) until bud formation. **(b)** Timing between the first Sec3 polarization event and SPB separation was scored in WT cells (*n* = 190), *bem1 cdc24$^{lip}$* cells that displayed a stable pole (*n* = 43), or unstable pole (*n* = 18 for unstable pole that form a bud, *n* = 131 for unstable pole that remain unbudded). **(c)** Frequency of multinucleate cells that appear after 1-NAA treatment in *bem1 cdc24$^{lip}$*, *bem1 cdc24$^{lip}$ cdc28$^{cdk1}$ Y19F* and *bem1 cdc24$^{lip}$ swe1$^{wee1}$Δ* cells (*n* > 100 cells per time point). Values display mean +/- SD. Unpaired *t*-tests with Welch corrections were performed on A and B. A two-way ANOVA was performed in C. (** $P < 0.01$, *** $P < 0.001$, **** $P < 0.0001$).

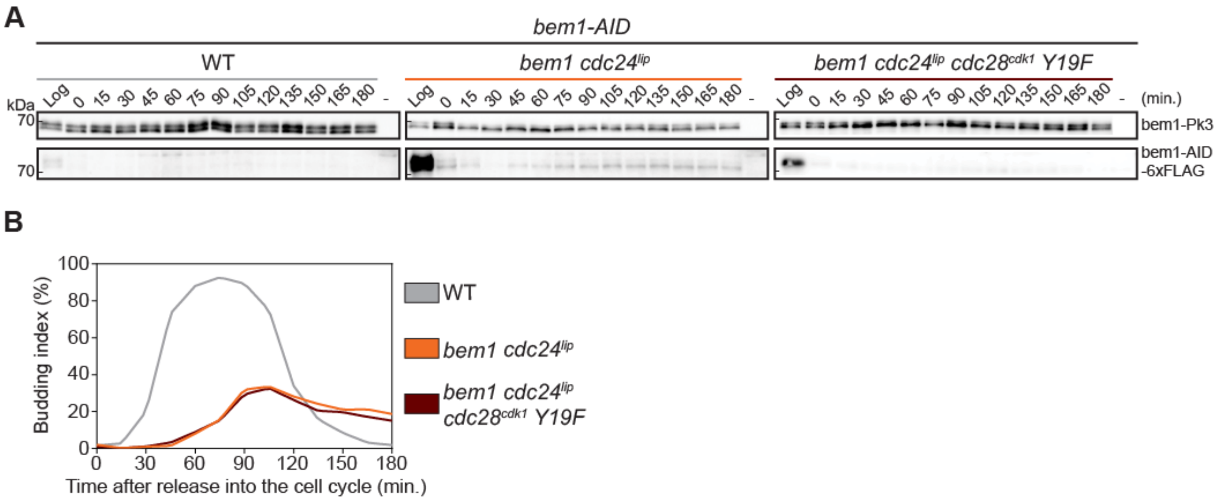


Supplementary Fig. 4

**Supplementary Fig. 4. Control Western blots and frequency of budding in *BEM1*, *bem1 cdc24$^{lip}$* and *bem1 cdc24$^{lip}$ cdc28$^{cdk1}$ Y19F* cells.**

The data in this Figure accompanies the cell cycle time-course shown in Figure 3. **(a)** Cells of the indicated genotypes were analyzed by Western blotting. **(b)** Frequency of budded cells after release into the cell cycle.

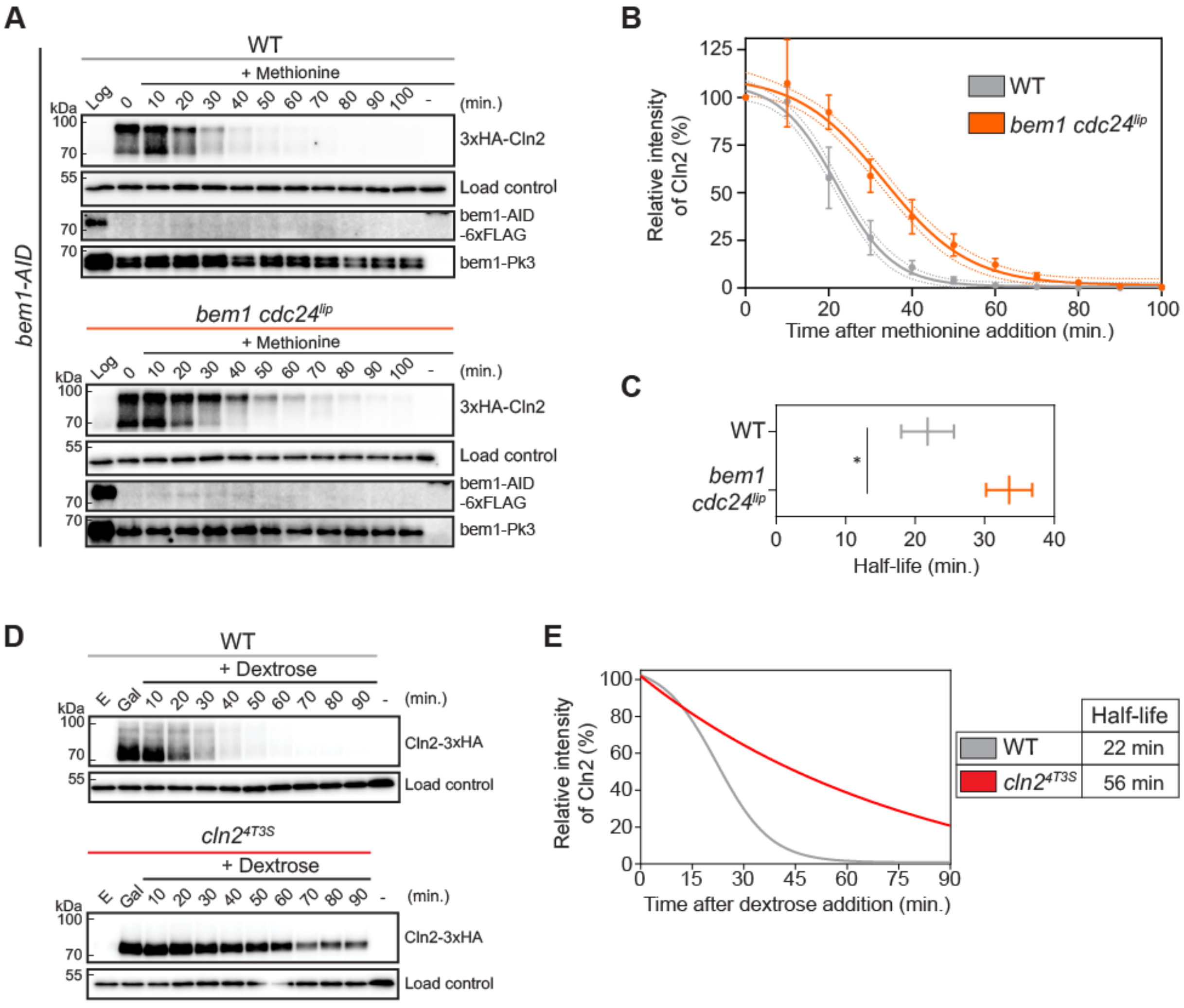


Supplementary Fig. 5

**Supplementary Fig. 5. The G1 cyclin Cln2 is stabilized in the *bem1 cdc24$^{lip}$* mutant.**

**(a)** 3xHA-Cln2, under control of the MET25 promoter, was expressed for 2 hours and then repressed by addition of methionine in WT cells or *bem1 cdc24$^{lip}$* mutant cells. Samples were removed at the indicated times. “Log”: uninduced sample grown in YPD + methionine. This experiment was repeated 5 times, with similar results. **(b)** Relative abundance of Cln2 protein was determined by nonlinear regression fitting (sigmoidal, $R^2$ of 0.98 for WT and 0.96 for *bem1 cdc24$^{lip}$*). Mean +/- SD are displayed at each time point. **(c)** Half-life of Cln2 protein in WT (n = 4) and *bem1 cdc24$^{lip}$* mutant cells (n = 5). Values display mean +/- SD. Mann-Whitney test was performed (*$P < 0.05$). **(d)** 3xHA-Cln2, under control of the GAL1 promoter, was expressed for 1 hour and then repressed by addition of dextrose in WT cells or *cln2$^{4T3S}$* mutant cells. Samples were removed at the indicated times. “E”: uninduced sample grown in YEP + ethanol + glycerol. “Gal”: induced sample after 1 hour in Galactose medium. **(e)** Relative abundance of

Cln2 protein was determined by nonlinear regression fitting (sigmoidal for WT cells ($R^2$ of 0.99), one phase decay for *cln2*$^{4T3S}$ cells ($R^2$ of 0.98)).

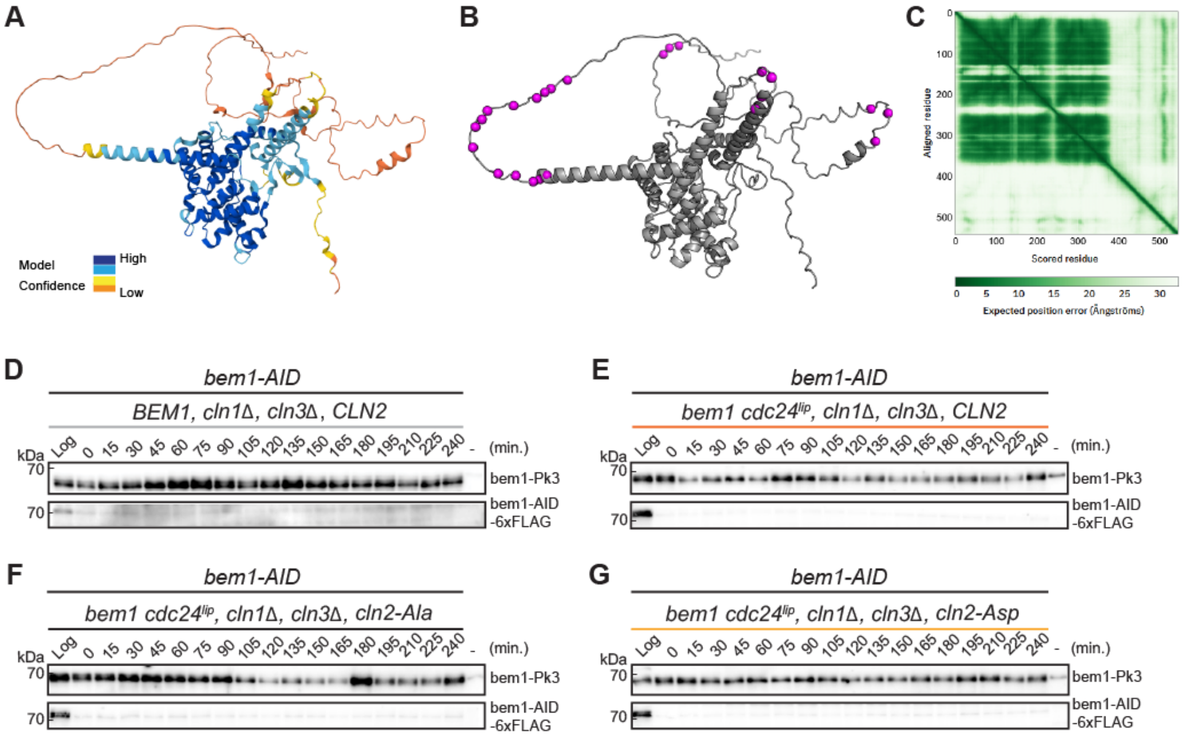


Supplementary Fig. 6

**Supplementary Fig. 6. G1 cyclin Cln2 structural model, expression of *cln2-Ala*/*cln2-Asp* phosphorylation mutants.**

**(a)** Alphafold model of Cln2 (https://alphafold.ebi.ac.uk/entry/P20438). **(b)** Alphafold model of Cln2 displaying the position of phosphorylation sites in pink. **(c)** Model confidence values. **(d-g)** Control Western blots for the time-courses in Figure 4.

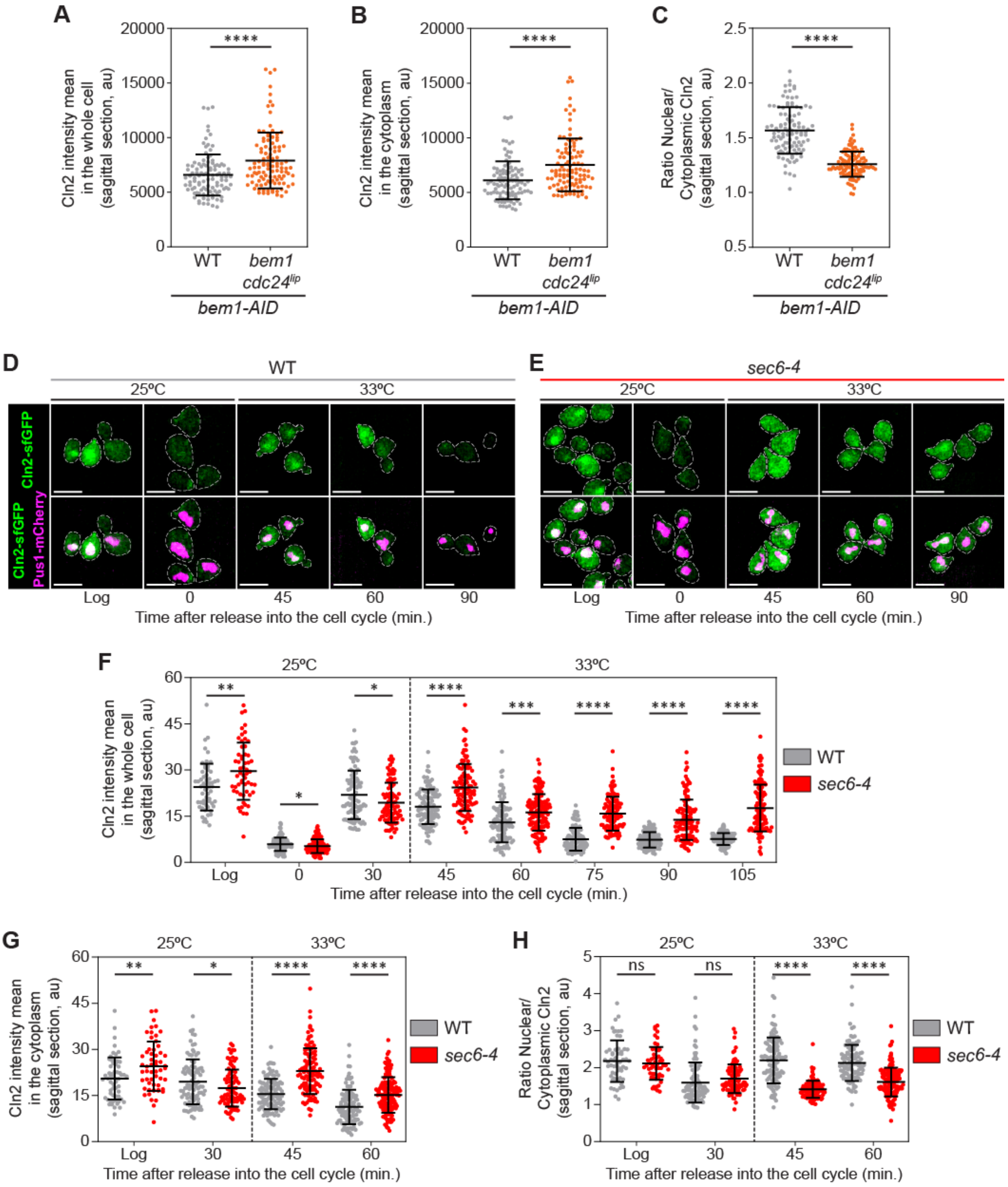


Supplementary Fig. 7

**Supplementary Fig. 7. A self-correcting mechanism that augments cytoplasmic G1 cyclin levels is triggered in response to polarity defects.**

**(a)** 1-NAA (0.5 mM) was added to log-phase cells for 5 hours to degrade bem1-AID in control (*n* = 104) or *bem1 cdc24*$^{lip}$ mutant cells (*n* = 108) and Cln2-sfGFP and Pus1-mRuby2 were imaged. Cln2 mean fluorescence intensity was extracted from a sagittal section through the cell, from which the nuclear and cytoplasmic signal was measured using the Pus1 signal to delineate the nucleoplasm. **(b)** Cln2 mean fluorescence intensity in the cytoplasm in WT cells (*n* = 104) and *bem1 cdc24$^{lip}$* mutant cells (*n* = 108). **(c)** The ratio of nuclear to cytoplasmic Cln2 fluorescence is displayed. **(d-e)** WT and *sec6-4* cells expressing Cln2-sfGFP (green) and Pus1-mCherry (magenta) were arrested in G1 with alpha-factor, then released into the cell cycle at 25ºC. 30 min after the release into the cell cycle, cells were shifted to the restrictive temperature (33ºC) and fixed every 15 minutes to measure Cln2 mean fluorescence intensity. Images are maximum-intensity projected z-stacks. Scale bars = 5 µm. **(f)** Quantification of Cln2 mean fluorescence intensity in the whole cell in WT cells and *sec6-4* mutant cells during the time course shown previously (*n* > 50 for Log time point, *n* > 100 cells for other time points). **(g)** Quantification of Cln2 mean fluorescence intensity in the cytoplasm in WT cells and *sec6-4* mutant cells during the time course shown previously (*n* > 50 for Log time point, *n* > 100 cells for other time points). **(h)** The ratio of nuclear to cytoplasmic Cln2 is displayed for WT cells and *sec6-4* mutant cells (*n* > 50 for Log time point, *n* > 100 cells for other time points). Values display mean +/- SD. Unpaired t-tests with Welch's correction were performed (* *P* < 0.05, ** *P* < 0.01, *** *P* < 0.001, **** *P*< 0.0001).

**Supplementary Table S1. Plasmids used in this study.**

| Name | Description |
|---|---|
| pDM364 (Addgene 20754 [67]) | pFA6a-*6xGLY-3xFLAG::kanMX6* |
| pDM443 | pRS303 |
| pDM526 | pFA6a-*pAID-6xFLAG::hphNT1* |
| pDM589 | *pTIR1::LEU2* |
| pDM996 | pFA6a-*GAGAGA-mRuby2::HIS3* |
| pDM1028 (Addgene 50632 [68]) | pFA6a-*pHIS3prom::Venus-Tub1+3'UTR::hphNT1* |
| pDM1045 | pUC19-*Pk3::LEU2* |
| pDM1046 | pFA6a-*sfGFP::kanMX6* |
| pDM1052 | pRS303 *pBEM1-bem1 bc-14E + px (K338M, K348A, R349A & R369A)-Pk3* |
| pDM1053 | pRS303 *BEM1-Pk3* |
| pDM1058 | pRS405 *CYC1promV3-PUS1-mRuby2-CYC1term* |
| pDM1064 | pFA6a-*GFP(S65T)::CaURA* |
| pDM1065 | pFA6a-*GFP(S65T)::natNT2* |
| pDM1069 (Addgene 44878 [69]) | pFA6a-*GAGAGA-yomKate2-CaURA3* |
| pDM1085 | pRS406 *CLN2-3xHA* |
| pDM1087 | pRS406 *cln2-3xHA* Ala (*S233A, S236A, S239A, T242A, T378A, S380A, T381A, S383A, S385A, S389A, S391A, T392A, S393A, S396A, S399A, S400A, S401A, S403A, T468A, S472A, T476A, S528A, S529A, S530A)* |
| pDM1090 | pRS406 *cln2-3xHA* Asp *(S233D, S236D, S239D, T242D, T378D, S380D, T381D, S383D, S385D, S389D, S391D, T392D, S393D, S396D, S399D, S400D, S401D, S403D, T468D, S472D, T476D, S528D, S529D, S530D)* |
| pDM1091 | pUC57 *cdc28prom-cdc28$^{cdk1}$Y19F-cdc28term::natNT2* |
| pDM1092 | pUC57 *cdc28prom-cdc28$^{cdk1}$Y19F-cdc28term::kanMX6* |
| pDM1133 | pFA6a- *mScarletI3::HIS3* |
| pDM1140 | pFA6a-*mScarletI3-mScarletI3::HIS3* |
| pDM1149 (Addgene 74636 [70]) | pFA6a-*mCherry::natNT2* |
| pYM21 [71] | pYM-*9MYC::natNT2* |
| pYM27 [71] | pYM-*eGFP::kanMX6* |
| pYM-N36 [71] | pYM-*MET25prom-3xHA::natNT2* |
| pFA6a-*natNT2* [71] | pFA6a-*natNT2* |
| [72] | pFA6a-*GFP(S65T)::kanMX6* |
| [72] | pFA6a-*GFP(S65T)::HIS3* |
| [72] | pFA6a-*3xHA::kanMX6* |

**Supplementary Table S2. Yeast strains used in this study.**

| DMY | Genotype |
|---|---|
| 255 | *MATa, his3Δ11, leu2Δ3,112, trp1Δ1, ura3Δ52, ade2Δ1, can1Δ100, GAL+, bar1, rga2Δ::kanMX6* |
| 552 | *MATa, his3Δ11, leu2Δ3,112, trp1Δ1, ura3Δ52, ade2Δ1, can1Δ100, GAL+, bar1, CLN2-3xHA::LEU2* |
| 808 | *MATa, his3Δ11, leu2Δ3,112, trp1Δ1, ura3Δ52, ade2Δ1, bar1, cln2Δ::HIS3, LEU2::pGAL1-CLN2-3xHA* |
| 809 | *MATa, his3Δ11, leu2Δ3,112, trp1Δ1, ura3Δ52, ade2Δ1, bar1, cln2Δ::HIS3, LEU2::pGAL1-cln2*$^{4T3S}$*-3xHA* |
| 996 | *MATa, his3Δ11, leu2Δ3, trp1Δ1, ura3Δ52, ade2Δ1, can1Δ100, GAL+,*<br>*HA-cdc28-T18A-Y19F:URA3* |
| 1054 | *MATa, his3Δ11, leu2Δ3,112, trp1Δ1, ura3Δ52, ade2Δ1, can1Δ100, GAL+, bar1, sec6-4::HIS3, CLN2-3xHA::LEU2* |
| 2105 | *MATa, his3Δ1, leu2Δ0, met15Δ0, ura3Δ0* |
| 2179 | *MATa, his3Δ1, leu2Δ0, met15Δ0, ura3Δ0, bem1Δ::CaURA3* |
| 2199 | *MATa, his3Δ1, leu2Δ0, met15Δ0, ura3Δ0, bem1 px (K338M, K348A, R349A & R369A)* |
| 2268 | *MATa, his3Δ1, leu2Δ0, met15Δ0, ura3Δ0, cdc24 K513A-3xHA::URA3* |
| 2472 | *MATa, his3Δ1, leu2Δ0, met15Δ0, ura3Δ0, bem1 bc-14E* |
| 2523 | *MATa, his3Δ1, leu2Δ0, met15Δ0, ura3Δ0, CDC24-3xHA::URA3, bem1 bc-14E + px (K338M, K348A, R349A & R369A)* |
| 2524 | *MATa, his3Δ1, leu2Δ0, met15Δ0, ura3Δ0, cdc24 K513A-3xHA::URA3, bem1 bc-14E + px (K338M, K348A, R349A & R369A)* |
| 2578 | *MATa, his3Δ1, leu2Δ0, met15Δ0, ura3Δ0, PUS1-GFP::HIS3* |
| 2591 | *MATa, his3Δ1, leu2Δ0, met15Δ0, ura3Δ0, cdc24 K513A-3xHA::URA3, bem1 bc-14E + px (K338M, K348A, R349A, R369A), PUS1-GFP::HIS3* |
| 2718 | *MATa, his3Δ1, leu2Δ0, met15Δ0, ura3Δ0, swe1Δ::kanMX6* |
| 2722 | *MATa, his3Δ1, leu2Δ0, met15Δ0, ura3Δ0, clb2Δ::kanMX6* |
| 2803 | *MATa, his3Δ1, leu2Δ0, met15Δ0, ura3Δ0, PUS1-mRuby2::HIS3, WHI5-sfGFP::kanMX6* |
| 2806 | *MATa, his3Δ1, leu2Δ0, met15Δ0, ura3Δ0, cdc24 K513A-3xHA ::URA3, bem1 bc-14E+px (K338M, K348A, R349A, R369A), PUS1-mRuby2::HIS3, WHI5-sfGFP::kanMX6* |
| 2910 | *MATa, his3Δ1, leu2Δ0, met15Δ0, ura3Δ0, SIC1-3xFLAG::kanMX6, CLB5-TAP::HIS3, CLN2-Pk3::LEU2* |
| 2912 | *MATa, his3Δ1, leu2Δ0, met15Δ0, ura3Δ0, cdc24 K513A-3xHA::URA3, bem1 bc-14E + px (K338M, K348A, R349A & R369A), SIC1-3xFLAG::kanMX6, CLB5-TAP::HIS3, CLN2-Pk3::LEU2* |
| 2957 | *MATa, his3Δ1, leu2Δ0, met15Δ0, ura3Δ0, cdc24 K513A-3xHA::URA3, leu2::LEU2::TIR1, bem1-AID-6xFLAG::hphNT1, his3::HIS3* |
| 2958 | *MATa, his3Δ1, leu2Δ0, met15Δ0, ura3Δ0, cdc24 K513A-3xHA::URA3, leu2::LEU2::TIR1, bem1-AID-6xFLAG::hphNT1, his3::BEM1-Pk3::HIS3* |
| 2960 | *MATa, his3Δ1, leu2Δ0, met15Δ0, ura3Δ0, cdc24 K513A-3xHA::URA3, leu2::LEU2::TIR1, bem1-AID-6xFLAG::hphNT1, his3::pBEM1-bem1 bc-14E + px (K338M, K348A, R349A & R369A)-Pk3::HIS3* |

| | |
|---|---|
| 2962 | *MATa, his3Δ1, leu2Δ0, met15Δ0, ura3Δ0, cdc24 K513A-3xHA::URA3, leu2::LEU2::TIR1, bem1-AID-6xFLAG::hphNT1, PUS1-mRuby2::kanMX6, his::HIS3* |
| 2963 | *MATa, his3Δ1, leu2Δ0, met15Δ0, ura3Δ0, cdc24 K513A-3xHA::URA3, leu2::LEU2::TIR1, bem1-AID-6xFLAG::hphNT1, PUS1-mRuby2::kanMX6, his::BEM1-Pk3::HIS3* |
| 2966 | *MATa, his3Δ1, leu2Δ0, met15Δ0, ura3Δ0, cdc24 K513A-3xHA::URA3, leu2::LEU2::TIR1, bem1-AID-6xFLAG::hphNT1, PUS1-mRuby2::kanMX6, his::pBEM1-bem1 bc-14E + px (K338M, K348A, R349A & R369A)-Pk3::HIS3* |
| 2978 | *MATa, his3Δ1, leu2Δ0, met15Δ0, ura3Δ0, Venus-TUB1::hphNT1, LEU2::Cyc1p-PUS1-mRuby2-Cyc1t, SPC42-eGFP::kanMX6* |
| 2980 | *MATa, his3Δ1, leu2Δ0, met15Δ0, ura3Δ0, cdc24 K513A-3xHA::URA3, bem1 bc-14E + px (K338M, K348A, R349A & R369A), Venus-TUB1::hphNT1, LEU2::Cyc1p-PUS1-mRuby2-Cyc1t, SPC42-eGFP::kanMX6* |
| 2996 | *MATa, his3Δ1, leu2Δ0, met15Δ0, ura3Δ0, cdc24 K513A, leu2::LEU2::TIR1, bem1-AID-6xFLAG::hphNT1, his::BEM1-Pk3::HIS3* |
| 2998 | *MATa, his3Δ1, leu2Δ0, met15Δ0, ura3Δ0, cdc24 K513A, leu2::LEU2::TIR1, bem1-AID-6xFLAG::hphNT1, his3::pBEM1-bem1 bc-14E + px (K338M, K348A, R349A & R369A)-Pk3::HIS3* |
| 3042 | *MATa, his3Δ1, leu2Δ0, met15Δ0, ura3Δ0, SEC3-GFP::kanMX6* |
| 3044 | *MATa, his3Δ1, leu2Δ0, met15Δ0, ura3Δ0, cdc24 K513A-3xHA::URA3, bem1 bc-14E + px (K338M, K348A, R349A & R369A), SEC3-GFP::kanMX6* |
| 3071 | *MATa, his3Δ1, leu2Δ0, met15Δ0, ura3Δ0, cdc24 K513A, leu2::LEU2::TIR1, bem1-AID-6xFLAG::hphNT1, his3::pBEM1-bem1 bc-14E + px (K338M, K348A, R349A & R369A)-Pk3::HIS3, CLN2-3HA::kanMX6* |
| 3081 | *MATa, his3Δ1, leu2Δ0, met15Δ0, ura3Δ0, cdc24 K513A, leu2::LEU2::TIR1, bem1-AID-6xFLAG::hphNT1, his::BEM1-Pk3::HIS3, CLN2-3HA::kanMX6, LTE1-9MYC::natNT2* |
| 3083 | *MATa, his3Δ1, leu2Δ0, met15Δ0, ura3Δ0, cdc24 K513A, leu2::LEU2::TIR1, bem1-AID-6xFLAG::hphNT1, his3::pBEM1-bem1 bc-14E + px (K338M, K348A, R349A & R369A)-Pk3::HIS3, CLN2-3HA::kanMX6, LTE1-9MYC::natNT2* |
| 3145 | *MATa, his3Δ1, leu2Δ0, met15Δ0, ura3Δ0, cdc24 K513A-3xHA::URA3, bem1 bc-14E + px (K338M, K348A, R349A & R369A), Venus-TUB1::hphNT1, LEU2::Cyc1p-PUS1-mRuby2-Cyc1t, SEC3-GFP::kanMX6* |
| 3176 | *MATa, his3Δ1, leu2Δ0, met15Δ0, ura3Δ0, cdc24 K513A, leu2::LEU2::TIR1, bem1-AID-6xFLAG::hphNT1, his3::pBEM1-bem1 bc-14E + px (K338M, K348A, R349A & R369A)-Pk3::HIS3, URA3::CLN2-3xHA, cln1Δ::kanMX6, cln3Δ::natNT2* |
| 3178 | *MATa, his3Δ1, leu2Δ0, met15Δ0, ura3Δ0, cdc24 K513A, leu2::LEU2::TIR1, bem1-AID-6xFLAG::hphNT1, his3::pBEM1-bem1 bc-14E + px (K338M, K348A, R349A & R369A)-Pk3::HIS3, URA3::cln2-3xHA (S233A, S236A, S239A, T242A, T378A, S380A, T381A, S383A, S385A, S389A, S391A, T392A, S393A, S396A, S399A, S400A, S401A, S403A, T468A, S472A, T476A, S528A, S529A, S530A), cln1Δ::kanMX6, cln3Δ::natNT2* |
| 3180 | *MATa, his3Δ1, leu2Δ0, met15Δ0, ura3Δ0, cdc24 K513A, leu2::LEU2::TIR1, bem1-AID-6xFLAG::hphNT1, his3::pBEM1-bem1 bc-14E + px (K338M, K348A, R349A & R369A)-Pk3::HIS3, URA3::cln2-3xHA (S233D, S236D, S239D, T242D, T378D, S380D, T381D, S383D, S385D, S389D, S391D, T392D, S393D, S396D, S399D, S400D, S401D, S403D, T468D, S472D, T476D, S528D, S529D, S530D), cln1Δ::kanMX6, cln3Δ::natNT2* |

| | |
|---|---|
| 3185 | *MATa, his3Δ1, leu2Δ0, met15Δ0, ura3Δ0, Venus-TUB1::hphNT1, LEU2::Cyc1p-PUS1-mRuby2-Cyc1t, SEC3-GFP::kanMX6* |
| 3202 | *MATa, his3Δ1, leu2Δ0, met15Δ0, ura3Δ0, cdc24 K513A, leu2::LEU2::TIR1, bem1-AID-6xFLAG::hphNT1, his::BEM1-Pk3::HIS3, URA3::CLN2-3xHA, cln1Δ::kanMX6, cln3Δ::natNT2* |
| 3206 | *MATa, his3Δ, leu2Δ0, met15Δ0, ura3Δ0, cdc24 K513A, leu2::LEU2::TIR1, bem1-AID-6xFLAG::hphNT1, his::BEM1-Pk3::HIS3, SEC3-GFP::KanMX6, SPC42-yomKate2-CaURA3* |
| 3207 | *MATa, his3Δ1, leu2Δ0, met15Δ0, ura3Δ0, cdc24 K513A, leu2::LEU2::TIR1, bem1-AID-6xFLAG::hphNT1, his3::pBEM1-bem1 bc-14E + px (K338M, K348A, R349A & R369A)-Pk3::HIS3, SEC3-GFP::KanMX6, SPC42-yomKate2-CaURA3* |
| 3230 | *MATa, his3Δ1, leu2Δ0, met15Δ0, ura3Δ0, cdc24 K513A, leu2::LEU2::TIR1, bem1-AID-6xFLAG::hphNT1, his3::pBEM1-bem1 bc-14E + px (K338M, K348A, R349A & R369A)-Pk3::HIS3, PUS1-GFP::CaURA* |
| 3234 | *MATa, his3Δ1, leu2Δ0, met15Δ0, ura3Δ0, cdc24 K513A, leu2::LEU2::TIR1, bem1-AID-6xFLAG::hphNT1, his3::pBEM1-bem1 bc-14E + px (K338M, K348A, R349A & R369A)-Pk3::HIS3, swe1Δ::kanMX6, PUS1-GFP::CaURA* |
| 3326 | *MATa, his3Δ1, leu2Δ0, met15Δ0, ura3Δ0, cdc24 K513A, leu2::LEU2::TIR1, bem1-AID-6xFLAG::hphNT1, his3::pBEM1-bem1 bc-14E + px (K338M, K348A, R349A & R369A)-Pk3::HIS3, cdc28::cdc28prom-cdc28 Y19F::kanMX6-cdc28term, PUS1-GFP::CaURA* |
| 3333 | *MATa, his3Δ1, leu2Δ0, met15Δ0, ura3Δ0, cdc24 K513A, leu2::LEU2::TIR1, bem1-AID-6xFLAG::hphNT1, his3::pBEM1-bem1 bc-14E + px (K338M, K348A, R349A & R369A)-Pk3::HIS3, cdc28::cdc28prom-cdc28 Y19F::kanMX6-cdc28term, URA3::CLN2-3xHA* |
| 3338 | *MATa, his3Δ1, leu2Δ0, met15Δ0, ura3Δ0, cdc24 K513A, leu2::LEU2::TIR1, bem1-AID-6xFLAG::hphNT1, his3::pBEM1-bem1 bc-14E + px (K338M, K348A, R349A & R369A)-Pk3::HIS3, cdc28::cdc28prom-cdc28 Y19F::kanMX6-cdc28term, SPC42-yomKate2-CaURA3, SEC3-GFP::natNT2* |
| 3350 | *MATa, his3Δ1, leu2Δ0, met15Δ0, ura3Δ0, CLB5-sfGFP::kanMX6, CLB2-mScarletI3-mScarletI3::HIS3* |
| 3352 | *MATa, his3Δ1, leu2Δ0, met15Δ0, ura3Δ0, cdc24 K513A-3xHA::URA3, bem1 bc-14E + px (K338M, K348A, R349A & R369A), CLB5-sfGFP::kanMX6, CLB2-mScarletI3-mScarletI3::HIS3* |
| 3370 | *MATa, his3Δ1, leu2Δ0, met15Δ0, ura3Δ0, cdc24 K513A, leu2::LEU2::TIR1, bem1-AID-6xFLAG::hphNT1, his3::pBEM1-bem1 bc-14E + px (K338M, K348A, R349A & R369A)-Pk3::HIS3, URA3::CLN2-3xHA, cln1Δ::kanMX6, SPC42-GFP::natNT2* |
| 3372 | *MATa, his3Δ1, leu2Δ0, met15Δ0, ura3Δ0, cdc24 K513A, leu2::LEU2::TIR1, bem1-AID-6xFLAG::hphNT1, his3::pBEM1-bem1 bc-14E + px (K338M, K348A, R349A & R369A)-Pk3::HIS3, URA3::cln2-3xHA (S233A, S236A, S239A, T242A, T378A, S380A, T381A, S383A, S385A, S389A, S391A, T392A, S393A, S396A, S399A, S400A, S401A, S403A, T468A, S472A, T476A, S528A, S529A, S530A), cln1Δ::kanMX6, SPC42-GFP::natNT2* |
| 3374 | *MATa, his3Δ1, leu2Δ0, met15Δ0, ura3Δ0, cdc24 K513A, leu2::LEU2::TIR1, bem1-AID-6xFLAG::hphNT1, his3::pBEM1-bem1 bc-14E + px (K338M, K348A, R349A & R369A)-Pk3::HIS3, URA3::cln2-3xHA (S233D, S236D, S239D, T242D, T378D, S380D, T381D, S383D, S385D, S389D, S391D, T392D, S393D, S396D, S399D, S400D, S401D, S403D, T468D, S472D, T476D, S528D, S529D, S530D), cln1Δ::kanMX6, SPC42-GFP::natNT2* |
| 3382 | *MATa, his3Δ1, leu2Δ0, ura3Δ0, bnr1Δ::kanMX6, bni1-FH2::HIS3, URA3::CLN2-3xHA* |

| | |
|---|---|
| 3443 | *MATa, his3Δ1, leu2Δ0, met15Δ0, ura3Δ0, cdc24 K513A, leu2::LEU2::TIR1, bem1-AID-6xFLAG::hphNT1, his::BEM1-Pk3::HIS3, CLN2-sfGFP-CLN2term::kanMX6, PUS1-mCherry::natNT2* |
| 3445 | *MATa, his3Δ1, leu2Δ0, met15Δ0, ura3Δ0, cdc24 K513A, leu2::LEU2::TIR1, bem1-AID-6xFLAG::hphNT1, his3::pBEM1-bem1 bc-14E + px (K338M, K348A, R349A & R369A)-Pk3::HIS3, CLN2-sfGFP-CLN2term::kanMX6, PUS1-mCherry::natNT2* |
| 3461 | *MATa, his3Δ11, leu2Δ3,112, trp1Δ1, ura3Δ52, ade2Δ1, can1Δ100, GAL+, bar1, sec6-4::HIS3, CLN2-3xHA::LEU2, CLN2-sfGFP-CLN2term::kanMX6, PUS1-mCherry::natNT2* |
| 3474 | *MATa, his3Δ1, leu2Δ0, met15Δ0, ura3Δ0, cdc24 K513A, leu2::LEU2::TIR1, bem1-AID-6xFLAG::hphNT1, his::BEM1-Pk3::HIS3, natNT2::MET25prom-3xHA-CLN2* |
| 3477 | *MATa, his3Δ1, leu2Δ0, met15Δ0, ura3Δ0, cdc24 K513A, leu2::LEU2::TIR1, bem1-AID-6xFLAG::hphNT1, his3::pBEM1-bem1 bc-14E + px (K338M, K348A, R349A & R369A)-Pk3::HIS3, natNT2::MET25prom-3xHA-CLN2* |
| 3482 | *MATa, his3Δ1, leu2Δ0, met15Δ0, ura3Δ0, cdc24 K513A, leu2::LEU2::TIR1, bem1-AID-6xFLAG::hphNT1, his3::pBEM1-bem1 bc-14E + px (K338M, K348A, R349A & R369A)-Pk3::HIS3, URA3::cln2-Ala (S233A, S236A, S239A, T242A, T378A, S380A, T381A, S383A, S385A, S389A, S391A, T392A, S393A, S396A, S399A, S400A, S401A, S403A, T468A, S472A, T476A, S528A, S529A, S530A)-sfGFP-CLN2term::kanMX6, PUS1-mCherry::natNT2* |
| 3484 | *MATa, his3Δ1, leu2Δ0, met15Δ0, ura3Δ0, cdc24 K513A, leu2::LEU2::TIR1, bem1-AID-6xFLAG::hphNT1, his3::pBEM1-bem1 bc-14E + px (K338M, K348A, R349A & R369A)-Pk3::HIS3, URA3::cln2-Asp (S233D, S236D, S239D, T242D, T378D, S380D, T381D, S383D, S385D, S389D, S391D, T392D, S393D, S396D, S399D, S400D, S401D, S403D, T468D, S472D, T476D, S528D, S529D, S530D)-sfGFP-CLN2term::kanMX6, PUS1-mCherry::natNT2* |

**Supplementary Movie 1. Sec3-GFP (exocyst) dynamics in a WT control cell.**

Images of WT cells expressing Sec3-GFP (green) acquired by time-lapse confocal spinning disk microscopy. Images are maximum-intensity projected z-stacks where frames were acquired every 3 minutes. Time is displayed in hours/minutes. The white arrow shows Sec3 polarization that delineates a stable polarity axis. This arrow turns red after the emergence of the bud. Scale bar = 5 µm. The video corresponds to Fig. 1a (upper panel).

**Supplementary Movie 2. Sec3-GFP (exocyst) dynamics in a *bem1 cdc24$^{lip}$* mutant cell displaying polarity establishment and maintenance defects.**

Images of *bem1 cdc24$^{lip}$* mutant cells expressing Sec3-GFP (green) acquired by time-lapse confocal spinning disk microscopy. Images are maximum-intensity projected z-stacks where frames were taken every 3 minutes. Time is displayed in hours/minutes. The white arrows show two unsuccessful Sec3 polarization attempts, before a final successful attempt that results in bud emergence (white arrow turns red). Scale bar = 5 µm. The video corresponds to Fig. 1a (lower panel).

**Supplementary Movie 3. The expressions of S- and M- phase cyclins are confined to brief temporal windows in WT cells.**

Images of WT cells expressing Clb5-sfGFP (S-phase cyclin, cyan) and Clb2-2xmScarletl3 (M-phase cyclin, magenta) acquired by time-lapse confocal spinning disk microscopy. Images are maximum-intensity projected z-stacks where frames were taken every 5 minutes. Time is displayed in hours/minutes. Colocalization of Clb5 and Clb2 is shown in white. Scale bar = 5 µm. The video corresponds to Fig. 1h (upper panel).

**Supplementary Movie 4. S- and M- phase cyclin expression is prolonged in *bem1 cdc24$^{lip}$* mutant cells.**

Images of *bem1 cdc24$^{lip}$* mutant cells expressing Clb5-sfGFP (S-phase cyclin, cyan) and Clb2-2xmScarletl3 (M-phase cyclin, magenta) acquired by time-lapse confocal spinning disk microscopy. Images are maximum-intensity projected z-stacks. Frames were acquired every 5 minutes and time is displayed in hours/minutes. Colocalization of Clb5 and Clb2 is shown in white. Scale bar = 5 µm. The cell on the left corresponds to Fig. 1h (middle panel), whereas the cell on the right shows an unbudded cell that displayed prolonged Clb2 expression.

**Supplementary Movie 5. Unbudded *bem1 cdc24$^{lip}$* mutant cells exhibit a mitotic delay, and become multinucleate if a daughter cell is not generated.**

Images of *bem1 cdc24$^{lip}$* mutant cells expressing Clb5-sfGFP (S-phase cyclin, cyan) and Clb2-2xmScarletl3 (M-phase cyclin, magenta) acquired by time-lapse confocal spinning disk microscopy. Images are maximum-intensity projected z-stacks where frames were acquired every 5 minutes. Time is displayed in hours/minutes. Colocalization of Clb5 and Clb2 is shown in white. Scale bar = 5 µm. The cell corresponds to Fig. 1h (lower panel).

**Supplementary Movie 6. Misordering of cell cycle events and the appearance of multinucleate cells.**

Images of a WT cell (left) and a *bem1 cdc24$^{lip}$* mutant cell (right) expressing Venus-Tub1 (microtubules, white), Spc42-eGFP (SPBs, white) and Pus1-mRuby2 (nucleoplasm, red) acquired by time-lapse confocal spinning disk microscopy. Images are maximum-intensity projected z-stacks where frames were taken every 2 minutes. Time is displayed in hours/minutes. Scale bar = 5 µm. In contrast to WT cells, where ordering of cell cycle events is invariant, mitotic spindle formation and nuclear division can occur before bud emergence in *bem1 cdc24$^{lip}$* mutant cells. As a consequence, these types of cells become multinucleate.

**Supplementary Movie 7. The time from SPB separation to anaphase onset is reduced in *bem1 cdc24$^{lip}$ cln2-Ala*, while being delayed in *bem1 cdc24$^{lip}$ cln2-Asp* mutant cells.**

Images of *bem1 cdc24$^{lip}$* (**A**), *bem1 cdc24$^{lip}$ cln2-Ala* (**B**) and *bem1 cdc24$^{lip}$ cln2-Asp* (**C**) mutant cells in the AID system expressing Spc42-GFP (SPBs, white) acquired by time-lapse confocal spinning disk microscopy. Images are maximum-intensity projected z-stacks where images were acquired every 5 minutes. Time is displayed in hours/minutes. Scale bar = 5 µm. The cells were synchronized in G1 phase, bem1-AID was degraded after auxin addition and cells were released into the cell cycle. The video corresponds to Fig. 4g.

**Supplementary Movie 8. The G1 cyclin Cln2 prominently localizes to the nucleus in WT cells.**

Images of WT cells in the AID system expressing Cln2-sfGFP (G1-phase cyclin, green) and Pus1-mCherry (nucleoplasm, magenta) acquired by time-lapse confocal spinning disk microscopy. Images are maximum-intensity projected z-stacks where frames were taken every 10 minutes. Time is displayed in hours/minutes. Scale bar = 5 µm. Colocalization of Cln2 and Pus1 is shown in white. The cells were synchronized in G1 phase, bem1-AID was degraded after auxin addition and cells were released into the cell cycle. The video corresponds to Fig. 5a (upper panel).

**Supplementary Movie 9. Prominent cytoplasmic localization of Cln2-sfGFP in *bem1 cdc24^lip^* mutant cells which is rapidly degraded after bud emergence.**

Images of *bem1 cdc24^lip^* mutant cells in the AID system expressing Cln2-sfGFP (G1-phase cyclin, green) and Pus1-mCherry (nucleoplasm, magenta) acquired by time-lapse confocal spinning disk microscopy. Images are maximum-intensity projected z-stacks where frames were acquired every 10 minutes. Time is displayed in hours/minutes. Scale bar = 5 µm. Colocalization of Cln2 and Pus1 is shown in white. The cells were synchronized in G1 phase, bem1-AID was degraded after auxin addition and cells were released into the cell cycle. The video corresponds to Fig. 5a (middle panel).

**Supplementary Movie 10. Unbudded *bem1 cdc24^lip^* mutant cells display enrichment and stabilization of cytoplasmic Cln2.**

Images of *bem1 cdc24^lip^* mutant cells in the AID system expressing Cln2-sfGFP (G1-phase cyclin, green) and Pus1-mCherry (nucleoplasm, magenta) acquired by time-lapse confocal spinning disk microscopy. Images are maximum-intensity projected z-stacks where frames were acquired every 10 minutes. Time is displayed in hours/minutes. Scale bar = 5 µm. Colocalization of Cln2 and Pus1 is shown in white. The cells were synchronized in G1 phase, bem1-AID was degraded after auxin addition and cells were released into the cell cycle. The video corresponds to Fig. 5a (lower panel).

**Supplementary Movie 11. The G1 cyclin Cln2 prominently localizes to the nucleus in unbudded *bem1 cdc24$^{lip}$ cln2-Ala* mutant cells.**

Images of *bem1 cdc24$^{lip}$ cln2-Ala* mutant cells in the AID system expressing Cln2-sfGFP (G1-phase cyclin, green) and Pus1-mCherry (nucleoplasm, magenta) acquired by time-lapse confocal spinning disk microscopy. Images are maximum-intensity projected z-stacks where frames were acquired every 10 minutes. Time is displayed in hours/minutes. Scale bar = 5 µm. Colocalization of Cln2 and Pus1 is shown in white. The cells were synchronized in G1 phase, bem1-AID was degraded after auxin addition and cells were released into the cell cycle. The video corresponds to Fig. 5c (upper panel).

**Supplementary Movie 12. Unbudded *bem1 cdc24$^{lip}$ cln2-Asp* mutant cells display enrichment and stabilization of cytoplasmic Cln2.**

Images of *bem1 cdc24$^{lip}$ cln2-Asp* mutant cells in the AID system expressing Cln2-sfGFP (G1-phase cyclin, green) and Pus1-mCherry (nucleoplasm, magenta) acquired by time-lapse confocal spinning disk microscopy. Images are maximum-intensity projected z-stacks where frames were acquired every 10 minutes. Time is displayed in hours/minutes. Scale bar = 5 µm. Colocalization of Cln2 and Pus1 is shown in white. The cells were synchronized in G1 phase, bem1-AID was degraded after auxin addition and cells were released into the cell cycle. The video corresponds to Fig. 5c (lower panel).
